# Orientation-invariant morphometry reveals a continuum of dendritic spine forms in layer II pyramidal neurons of the petavoxel human connectome

**DOI:** 10.64898/2026.06.25.734571

**Authors:** Miguel A. Zamora-Ursulo, Elias Manjarrez

**Author notes:** **Corresponding Author:Prof. Elias Manjarrez,** Institute of Physiology, Benemérita Universidad Autónoma de Puebla. 14 sur 6301, Col. San Manuel A.P. 406, C.P. 72570. Puebla, Pue., México. Tels.

## Abstract

A recent study (Manjarrez et al., 2026) showed that the classification of cortical dendritic spines into stubby, thin, and mushroom subtypes is unstable under rotation. That result criticizes the categorical scheme but leaves an open question. What is the actual structure of spine morphology once the viewing angle is controlled? Here we answer it. We analyzed 228 spines from layer II pyramidal neurons in the H01 nanometer-resolution reconstruction of human temporal cortex. We first quantified the source of instability. We found that rotating dendritic segments by 90 degrees about their axes shifted the apparent spine height and head width in opposite directions across the population, thereby confirming orientation-dependent measurement error. Furthermore, to obtain measurements free of this artifact, we developed the Spine Morphometry Hub (SMH), a 12-point anatomical landmark framework that characterizes each spine in all three orthogonal planes and extracts geometric, voxel-based, and mesh-based metrics. All morphometric distributions were unimodal and right-skewed. Density-based clustering assigned most spines to noise, and a Monte-Carlo test against a discrete two-type null model confirmed that this pattern is incompatible with categorical subtypes. We also confirmed that apical and basal spines were statistically indistinguishable. Unlike previous reports of a spine continuum, all based on orientation-dependent measurements, our framework removes the viewing-angle confound itself, so the continuum we observe cannot be attributed to a projection artifact. Hence, our framework will be useful to quantify dendritic-spine remodeling in neurological disorders, in which spine shape has long been observed but never measured against an orientation-invariant morphometric standard.

**Highlights:**

- Spine Morphometry Hub (SMH) measures spines free of viewing-angle error
- SMH was validated as an orientation-invariant morphometry framework
- Rotating dendrites by 90° shifts spine height and head width oppositely
- All morphometric distributions are unimodal and right-skewed, not categorical
- SMH could be used to quantify dendritic-spine remodeling in neurological disorders

## Introduction

Dendritic spines are actin-rich postsynaptic protrusions that receive the vast majority of excitatory glutamatergic input in the mammalian neocortex and constitute the primary structural basis of synaptic weight (Harris & Kater, 1994; Yuste & Bonhoeffer, 2001, 2004; Yuste, 2011; Alvarez & Sabatini, 2007). Their geometry is not merely anatomical because head volume scales with postsynaptic density area and AMPA receptor content, linking morphology directly to synaptic efficacy (Matsuzaki et al., 2001; Harris & Stevens, 1989; Lisman & Harris, 1993), and trans-synaptic adhesion complexes such as neuroligin-neurexin pairs stabilize contact geometry (Südhof, 2008). The neck modulates biochemical and electrical compartmentalization relative to the parent dendrite, filtering the diffusion of second messengers and attenuating depolarization (Koch & Zador, 1993; Yuste, 2013; Tonnesen et al., 2014; Araya et al., 2014; Cornejo et al., 2022).

Structural remodeling of spines underlies long-term potentiation and depression and is considered the morphological substrate of memory encoding (Kasai et al., 2010; Bosch & Hayashi, 2012; Yasuda et al., 2022). In vivo imaging has revealed that a fraction of spines persists for months while others turn over rapidly (Grutzendler et al., 2002; Trachtenberg et al., 2002), and this structural plasticity involves actin cytoskeleton reorganization (Yang & Liu, 2022), synapse-specific signaling cascades (Colgan et al., 2023; Harris et al., 2024), and mechanical coupling across the cleft (Kasai et al., 2023). Pathological alterations in spine density and geometry are among the most consistently reported anatomical findings in schizophrenia, Alzheimer’s disease, and neurodevelopmental disorders (Penzes et al., 2011; Glantz & Lewis, 2000; Glausier & Lewis, 2013; Terry et al., 1991). Therefore, establishing the three-dimensional geometric diversity of human cortical spines is a prerequisite for interpreting both normal synaptic function and disease-related structural remodeling.

The classical morphological classification of dendritic spines into stubby, thin, and mushroom subtypes, proposed by Peters and Kaiserman-Abramof (1970) based on Golgi-impregnated material examined in two-dimensional projections, has shaped the field for over five decades. This categorical scheme was operationally useful but was established without quantitative criteria and from projections inherently dependent on viewing angle, as demonstrated in a recent study (Manjarrez et al., 2026).

However, that demonstration established the problem but not its resolution. If categorical assignment is unstable because it depends on viewing angle, the underlying structure of spine morphology remains unknown until the angle is controlled. Here we take that step. Rather than classifying spines from a fixed viewpoint, we introduce an orientation-invariant landmark framework, as Manjarrez et al. 2026 propose. This landmark measures each spine in a frame intrinsic in its own geometry, and we ask whether discrete subtypes re-emerge once the viewing-angle artifact is removed.

This question does not arise in a vacuum. Seminal three-dimensional reconstructions using serial-section electron microscopy have reported that most spines in the mouse visual cortex (Arellano et al., 2007) and the rat hippocampus (Harris & Stevens, 1989) exhibit intermediate morphologies that resist unambiguous categorical assignment. Arellano et al. (2007) examined 144 fully electron-microscopy-reconstructed spines from mouse visual cortex layer II/III and explicitly concluded that no clear subgrouping could be detected. More recent ultrastructural analyses confirmed continuous distributions in rodent (Ofer et al., 2021) and human (Ofer et al., 2022) cortical spines, and systematic comparisons of spine dimensions across cortical regions and species have reinforced these findings (Benavides-Piccione et al., 2025).

Computational geometry approaches applied to structured illumination microscopy (Kashiwagi et al., 2019) and confocal reconstructions of over 7,000 human pyramidal cell spines (Luengo-Sanchez et al., 2018) likewise demonstrated that spine shape descriptors are distributed along continuous axes, prompting a formal reappraisal of the classification-versus-clusterization debate (Pchitskaya & Bezprozvanny, 2020). Despite this accumulating evidence, discrete categorical classification remains widespread, partly because tools enabling systematic orientation-invariant morphometry with explicit quality control and cross-method validation in human tissue have been lacking.

Beyond the descriptive question of whether spine shapes are discrete or continuous, this topology has direct functional consequences. Each combination of head volume and neck dimensions corresponds to a unique pairing of synaptic weight and biochemical compartmentalization (Matsuzaki et al., 2001; Koch & Zador, 1993; Tonnesen et al., 2014). If the distribution of these structural parameters is continuous, the computational diversity of excitatory synapses is also continuous. Two-photon imaging studies have established that changes in spine head volume are graded and proportional to the magnitude of the plasticity-inducing stimulus, rather than stepwise (Matsuzaki et al., 2004; Holtmaat & Svoboda, 2009). Spine necks respond to activity within minutes through myosin- and cofilin-dependent actin remodeling, changing compartmentalization independently of head size (Tonnesen et al., 2014; Araya et al., 2014). Therefore, understanding the true topology of morphological space in human tissue is a prerequisite for interpreting normal cortical function and pathological deviations.

Previous volume electron microscopy studies have provided near-isotropic, nanometer-scale resolution in three dimensions (Bock et al., 2011; Kasthuri et al., 2015; Motta et al., 2019; Shapson-Coe et al., 2024; Lichtman & Denk, 2011), and the public release of the H01 human connectome and the MICrONS mouse dataset (Turner et al., 2022; Loomba et al., 2022) has provided unprecedented resources for addressing questions regarding connectivity and structure at nanometer resolution.

Here we implement this framework as the Spine Morphometry Hub (SMH), a validated 12-point anatomical landmark system of orientation-invariant morphometry applied to dendritic spines from layer II pyramidal neurons in the H01 human connectome described above. Our findings establish a validated quantitative baseline for measuring pathological spine remodeling in neurological and psychiatric disorders.

## Methods

### Human connectome dataset

All morphometric analyses were performed on the H01 dataset, a publicly available nanometer-resolution volume electron microscopy reconstruction of approximately 1 mm³ block of human temporal cortex (Brodmann area 21, right hemisphere) from a 45-year-old female donor with no known neurological disease (Shapson-Coe et al., 2024). The tissue was acquired by serial-section transmission electron microscopy at 8 × 8 × 33 nm voxel resolution (x, y, z). Neuronal segmentation was performed by the H01 consortium using flood-filling networks (Januszewski et al., 2018), and triangulated surface meshes were generated at multiple levels of detail (LOD). All data access was via the CloudVolume Python interface (Van Rossum & Drake, 2009) using the segmentation source precomputed://gs://h01-release/data/20210601/c3 at LOD-0. No human subjects were recruited for this study; all analyses used the publicly released de-identified dataset.

### Neuron selection and dendritic compartment classification

Five layer II pyramidal neurons were selected for analysis (H01 segment IDs: 329164988, 329209038, 343679509, 329384517, 2103144214) on the basis of: (i) reconstruction of both apical and basal dendritic trees within the H01 volume, verified by visual inspection in Neuroglancer; (ii) absence of segmentation artefacts at the dendritic level; and (iii) sufficient spine density across multiple dendritic fragments to support statistical analysis. All five neurons were confirmed as layer II pyramidal cells by their characteristic morphology (pyramidal soma, apical dendrite extending toward the pial surface, basal dendritic skirt) and by depth within the H01 volume consistent with layer II of the middle temporal gyrus. All five neurons carry the label “C2” in the H01 dataset.

For each neuron, dendritic fragments were classified as apical or basal based on their trajectories relative to the soma: apical fragments extend toward the pial surface and are identifiable by their ascending, tapering morphology; basal fragments project laterally and inferiorly. The number of annotated fragments varied by neuron depending on the extent of dendritic reconstruction available within the imaged volume: three apical and three basal fragments for neurons 329164988 and 343679509; two apical and two basal for neurons 329209038 and 329384517; and one apical and one basal for neuron 2103144214, yielding 22 dendritic fragments in total (11 apical, 11 basal). Compartment labels were assigned at the fragment level before any spine annotations were examined, ensuring that morphometric measurements were obtained without reference to compartment assignment.

### Spine identification and annotation

Individual spines were identified in Neuroglancer as protrusions arising from the dendritic shaft with a discernible constriction at the base (neck) and an expanded terminal domain (head). Filopodia with no discernible head were included in the raw count. Branched spines were included if both branches were within the H01 volume. A total of 228 spines were annotated across the five neurons: neuron 329164988 (n = 74), 329209038 (n = 53), 343679509 (n = 43), 329384517 (n = 39), and 2103144214 (n = 19). All annotations were performed by a single experienced annotator (December 2025 to February 2026) following a 50-spine training phase and iterative refinement of the protocol.

### Spine Morphometry Hub: a 12-point anatomical landmark framework

The Spine Morphometry Hub (SMH) defines 12 anatomically constrained landmark points per spine in Neuroglancer and examines the spine simultaneously in all three orthogonal planes (XY, XZ, YZ). The landmarks are organized into three compartmental groups. Group 1 (insertion): P1_BASE, placed at the geometric center of the shaft–neck constriction, identified as the local minimum of the cross-sectional profile in transverse view. Group 2 (neck): two pairs of surface-constrained landmarks at the neck surface at approximately 25% (NECK_1A, NECK_1B) and 75% (NECK_2A, NECK_2B) of estimated neck length, placed on diametrically opposite sides of the neck in the plane orthogonal to the neck axis. Group 3 (head): P2_UNION at the neck–head junction; C_HEAD at the estimated geometric centroid of the head; TOP_HEAD at the most distal surface point; SIDE1_HEAD and SIDE2_HEAD at diametrically opposite lateral surface points; FRONT_HEAD and BACK_HEAD at diametrically opposite surface points orthogonal to the lateral pair. Points designated as surface-constrained are snapped to the nearest mesh vertex; internal points (P1_BASE, P2_UNION, C_HEAD) are positioned freely within the volume. Landmark coordinates were recorded in nanometers in the H01 global coordinate system and exported as CSV files.

### Geometric metrics (Block 1)

All geometric metrics were computed analytically from the 12 landmark coordinates using SMH.py (Python 3.8; NumPy, SciPy). Neck length: Euclidean distance from P1_BASE to P2_UNION. Neck diameters: d₂₅ = ||NECK_1A − NECK_1B|| and d₇₅ = ||NECK_2A − NECK_2B|| at the 25% and 75% neck positions; mean, minimum and maximum thereof. Neck taper ratio: d₇₅ / d₂₅ (values > 1 indicate convergence toward the head). Neck constriction index: d_min / d_mean. Neck volume (frustum): V = πh/3 · (r₂₅² + r₂₅r₇₅ + r₇₅²). Neck lateral area (frustum): A = π(r₂₅ + r₇₅) · g, where g = √(h² + (r₇₅ − r₂₅)²). Neck resistance proxy: L_neck / r_mean⁴ (Hagen–Poiseuille approximation). Head semi-axes: a = mean(||SIDE1 − C||, ||SIDE2 − C||), b = mean(||FRONT − C||, ||BACK − C||), c = ||TOP − C||. Head volume (ellipsoid): V = (4/3)πabc. Head equivalent diameter: 2(3V/4π)^(1/3). Head surface area (Knud–Thomsen approximation, error < 1.06%): A ≈ 4π[(a^p b^p + a^p c^p + b^p c^p)/3]^(1/p), p = 1.6075. Head aspect ratios: c/a and a/b. Spine total length: ||TOP_HEAD − P1_BASE||. Axes angle: angle between the neck vector (P1→P2) and the head vector (P2→TOP). Head/neck diameter ratio: head equivalent diameter / d_min. Spine curvature index: perpendicular deviation of P2 from the straight axis P1→TOP, normalized by total spine length.

### Voxel-based volumetry (Block 2)

Voxel-based volumes were computed from the H01 segmentation mask. For each spine, the mask was downloaded via CloudVolume within a bounding box encompassing all 12 landmarks plus a 1,500 nm margin. The binary mask (voxels matching the parent neuron segment ID) was isolated using a multi-step pipeline: (1) planar cut at P1_BASE perpendicular to the neck axis (P1→P2) to separate the spine from the dendritic shaft; (2) a curved tube filter following the anatomical waypoints of the spine (P1 → mid(NECK_1A,1B) → mid(NECK_2A,2B) → P2 → C_HEAD → TOP_HEAD) with locally interpolated radius (1.3× measured neck radius in the neck segments, 1.4× maximum head radius in the head segments; minimum 120–150 nm) to exclude shaft voxels captured by the watershed; (3) connected-component extraction seeded at C_HEAD, with KD-tree rescue if the seed voxel falls in background (tolerance 1.0 µm in physical distance); (4) removal of fragments < 5 voxels; and (5) separation into neck and head compartments by a planar cut at P2_UNION perpendicular to the neck axis. Voxel volume was computed as N_voxels × V_voxel, where V_voxel = 8 × 8 × 33 nm³ = 2.112 × 10⁻⁶ µm³.

Voxel-based metrics include: spine volume, head volume, neck volume, head percentage, neck/head volume ratio, and head PCA axes (semi-axes estimated as √5 × √λ from eigenvalue decomposition of the head voxel cloud covariance matrix, appropriate for uniformly distributed ellipsoidal point clouds), head anisotropy (1 − λ₃/λ₁), and head elongation (λ₁/λ₂).

### Mesh-based surface and volume metrics (Block 3)

Surface area and mesh-derived volumes were computed from the LOD-0 triangulated surface mesh downloaded via CloudVolume. The mesh processing pipeline comprised: (0) topological repair (clean, triangulate, consistent normal orientation via auto_orient_normals); (1) clipping at the P1_BASE plane (normal: P1→P2 axis) combined with the curved tube vertex filter described above to isolate the spine protrusion from the shaft; (2a) clipping at the P2_UNION plane to extract the head sub-mesh; (2b) the complementary clip to extract the neck sub-mesh.

Open boundaries introduced by planar clipping were closed using a stepped strategy: fan-cap triangulation from the boundary centroid; if unsuccessful, Delaunay 2D triangulation of boundary vertices; as a final fallback, fill_holes with iteratively increasing hole size. Volume was computed by the signed-tetrahedron (divergence) method: V = (1/6)|Σ v₀ · (v₁ × v₂)| (Bourne & Harris, 2008). Surface area was computed after Taubin smoothing (λ = 0.5, µ = −0.53, 15 iterations; Taubin, 1995) to correct staircase artifacts from voxelization while preserving volume (isotopic rescaling applied if volume change exceeded 5%). Sphericity was computed as ψ = π^(1/3)(6V)^(2/3)/A.

Mesh viability (mesh_viable = 1) required watertight meshes with positive volume for both head and neck sub-meshes, and mesh-to-geometric surface area ratios below sanity thresholds (neck mesh area < 6× geometric neck area; spine mesh area < 5× geometric total area) to exclude cases of inadvertent shaft capture.

### Soma distance estimation

Soma distance was estimated as the geodesic path length along the dendritic shaft surface from P1_BASE to the soma centroid, computed using the multi-path surface geodesic algorithm. The soma centroid was identified by visual inspection in Neuroglancer and confirmed by the characteristic pyramidal soma morphology in the electron microscopy volume.

### Quality control and data filtering

The primary analytical filter for the inclusion of dendritic spines in this study was vol_viable = 1, which required the conjunction of the following four conditions: (i) high measurement confidence (”alta”), defined as voxel–mesh head volume discrepancy < 20% and total volume ratio between 0.6 and 2.0; (ii) a finite voxel-based spine volume; (iii) absence of any quality-control flags (including RUIDO [head volume < 0.01 µm³], FUSION [spine volume > 2.0 µm³], DEMASIADO_LARGA [total length > 5.0 µm], MESH_INVALIDO, VOX_SEED_FAIL, SHAFT_VOX [voxel spine volume > 10× geometric total volume], and GEOM_ERROR); and (iv) voxel–mesh total volume discrepancy < 50%. A total of 228 spines passed this filter (vol_viable = 1; apical n = 115, basal n = 113). Hence, the final analytical dataset comprised n = 228 spines from five neurons: neuron 329164988 (n = 74; apical 36, basal 38), 329209038 (n = 53; apical 28, basal 25), 343679509 (n = 43; apical 22, basal 21), 329384517 (n = 39; apical 20, basal 19) and 2103144214 (n = 19; apical 9, basal 10).

### Variable selection for dimensionality reduction

For principal component analysis, a subset of seven morphometrically independent variables was selected from the full set of variables. Strongly correlated variable pairs (Spearman |ρ| ≥ 0.70) were identified from the full correlation matrix. From each redundant cluster, the single representative with the highest functional relevance to established biophysical models was retained: head volume [voxel] as the primary determinant of synaptic strength (Matsuzaki et al., 2001); head sphericity [mesh] as the geometric shape descriptor most directly linked to curvature analyses (Kashiwagi et al., 2019); neck length and neck diameter as the primary determinants of neck resistance (Koch & Zador, 1993); neck taper ratio as an independent shape descriptor of neck geometry; head/neck diameter ratio as the classical compartmentalization index (Harris et al., 1992); and axes angle as a three-dimensional geometric variable with no strong correlates in the head or neck domains (|ρ| < 0.30 with all other selected variables). This selection (Set-B) was designed to maximize the number of independent morphological dimensions while using the primary metric for each domain: voxel-based for volume, mesh-based for shape.

### Principal component analysis

The seven selected variables were standardized using RobustScaler (median subtraction, interquartile range scaling) to reduce sensitivity to outliers and reduced by PCA via full singular value decomposition. The number of retained components was the minimum required to explain ≥90% of total variance. Effective dimensionality was computed as D_eff = (Σλᵢ)² / Σλᵢ² (Del Giudice, 2021). Bootstrap 95% confidence intervals for variance explained were obtained from 250 stratified resamples.

### Density-based clustering

HDBSCAN (Hierarchical Density-Based Spatial Clustering of Applications with Noise; Campello et al., 2013; McInnes et al., 2017) was applied to the standardized seven-dimensional feature space (prior to PCA projection) with minimum cluster size = 7, minimum samples = 3, and excess-of-mass (EOM) cluster selection. HDBSCAN identifies regions of unusually high local density without requiring a predetermined number of clusters or assuming parametric cluster shapes; points that do not belong to any dense region are labeled as noise, indicating that they do not belong to any discrete cluster. The silhouette coefficient was computed on assigned (non-noise) points as a measure of cluster separation.

### Monte Carlo null hypothesis testing

To determine whether the high HDBSCAN noise rate observed in the real data reflects genuine biological continuity rather than algorithmic insensitivity, we performed a Monte Carlo simulation under the null hypothesis (H₀) that dendritic spines belong to a small number of discrete morphological types. The optimal number of discrete types under H₀ was determined by fitting Gaussian Mixture Models (GMMs) with k = 2 to 6 components to the real PCA-transformed data and selecting the model with the lowest Bayesian Information Criterion (BIC); this procedure selected k = 2 as the best-fitting discrete model.

For each of 1,000 independent iterations, a synthetic dataset of n = 228 points was drawn from two multivariate Gaussian distributions centered at the real HDBSCAN cluster centroids in the six-dimensional PCA space, with within-cluster standard deviation set to 30% of the empirical per-dimension standard deviation, producing distributions substantially more compact than any biologically realistic discrete taxonomy. HDBSCAN was applied to each synthetic dataset with identical parameters. The empirical p-value was defined as the proportion of simulations in which the H₀ noise rate was equal to or exceeded the observed real noise rate.

### Statistical analysis

Morphometric variables are described as mean ± standard deviation and median [interquartile range]. Normality was assessed by the Shapiro-Wilk test. Pairwise Spearman rank correlations were computed for all variable pairs; all tests were two-tailed at α = 0.05. Comparisons between apical and basal spines were performed using linear mixed-effects models (LMMs) fitted with restricted maximum likelihood (REML). Each morphometric variable was modeled as a function of dendritic compartment (apical versus basal; fixed effect) with neuron identity as a random intercept to account for the non-independence of spines sampled from the same neuron (n = 5 neurons, n = 228 spines).

Significance of the fixed effect was assessed via Wald z-tests. To control for multiple comparisons across 23 morphometric variables, p-values were adjusted using the Benjamini-Hochberg false discovery rate (FDR) procedure; adjusted p-values (q- values) < 0.05 were considered significant. Effect sizes are reported as Cohen’s d (pooled standard deviation). Intraclass correlation coefficients (ICC(1)) were computed per variable to quantify the proportion of variance attributable to neuron grouping. Mann–Whitney U tests are reported as sensitivity analyses (spine-level, ignoring clustering) for comparison with prior literature.

Compartment separability in the multivariate morphospace was assessed by logistic regression (L2 regularization, C = 1.0) trained on principal component scores with 5-fold stratified cross-validation (area under the ROC curve as the performance metric). Kolmogorov-Smirnov tests were used to compare score distributions between compartments along each principal component.

### Software and code availability

All analyses were performed in Python 3.8.10. Key libraries and versions: NumPy 1.24 (Harris et al., 2020), SciPy 1.11.1 (Virtanen et al., 2020), pandas 2.0 (McKinney, 2010), scikit-learn 1.3.0 (Pedregosa et al., 2011), PyVista 0.43 (Sullivan & Kaszynski, 2019), CloudVolume 8.x, HDBSCAN 0.8.33 (McInnes et al., 2017), statsmodels (for LMM). The SMH annotation and analysis pipeline (SMH.py) is available at [URL to be provided upon acceptance].

## Results

### 2D projections distort spine measurements

To quantify orientation-dependent measurement error, spine heights were measured before and after 90° rotation of the dendritic axis (n = 86 spines from manual 2D projections) (Fig. 1a-b). Spines were stratified into those whose apparent height increased (n = 43; from 1.07 ± 0.52 to 1.92 ± 0.74 µm; paired t-test, p < 0.001; Fig. 1c) and those whose apparent height decreased (n = 43; from 1.60 ± 0.66 to 1.00 ± 0.41 µm; p < 0.001; Fig. 1d). A Bland-Altman analysis confirmed no systematic directional bias at the group level (mean bias = +0.12 µm, p = 0.208), but the 95% limits of agreement spanned -1.64 to +1.89 µm (Fig. 1e), indicating biologically significant individual-spine error.

**Fig. 1.**
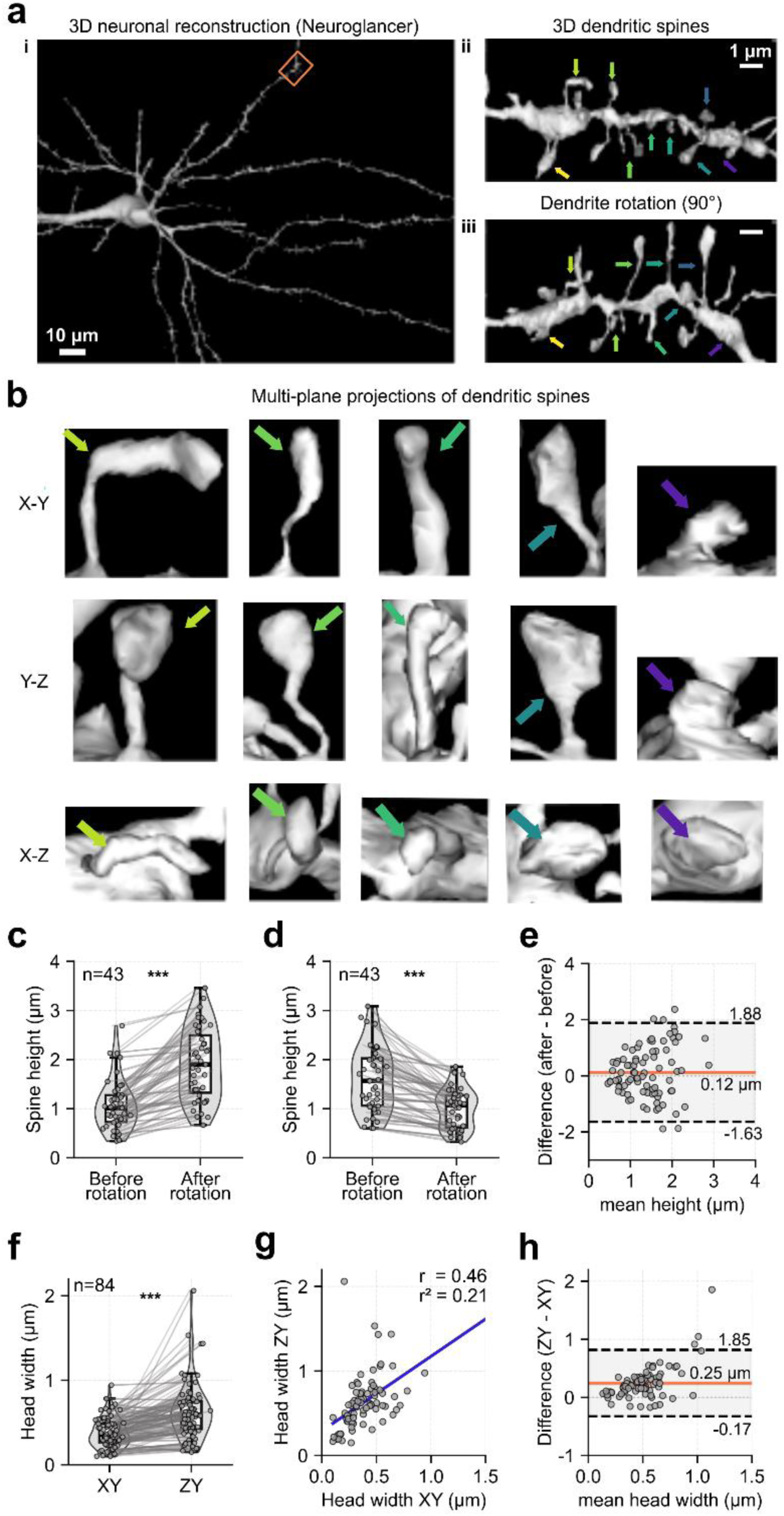
Two-dimensional projections introduce systematic measurement error in dendritic spine morphometry. **a**, Three-dimensional reconstruction of a layer II pyramidal neuron from the H01 dataset rendered in Neuroglancer. Panel I: whole-neuron view (scale bar, 10 µm); red rectangle delineates the analyzed segment. Panel ii: magnified view showing four representative spines (colored arrows; scale bar, 1 µm). Panel iii: the same segment after 90° rotation about the dendritic axis, revealing orientation-dependent changes in apparent spine profiles. **b**, Multi-plane orthogonal projections (X-Y, Y-Z, X-Z) of the four spines, illustrating that the same spine exhibits markedly different apparent morphology across projection planes. **c**, Paired comparison of spine height before and after 90° rotation for spines in which apparent height increased (n = 43). Mean height rose from 1.07 ± 0.52 µm to 1.92 ± 0.74 µm (paired t-test, p < 0.001). **d**, Complementary subgroup in which apparent height decreased (n = 43). Mean height fell from 1.60 ± 0.66 µm to 1.00 ± 0.41 µm (paired t-test, p < 0.001). **e**, Bland-Altman plot for spine height (n = 86). Mean bias = +0.12 µm (p = 0.208, one-sample t-test); 95% limits of agreement: -1.64 to +1.89 µm. **f**, Paired comparison of spine head width in X-Y versus Z-Y planes (n = 84; Wilcoxon signed-rank test, p < 0.001). **g**, Scatter plot of head width X-Y versus Z-Y (n = 84; Pearson r = 0.46). **h**, Bland-Altman plot for head width. Mean bias = +0.25 µm; limits of agreement: -0.17 to +1.81 µm.

Head width measured in X-Y versus Z-Y planes (n = 84) differed significantly (Wilcoxon signed-rank test, p < 0.001; Fig. 1f), with the Z-Y plane yielding systematically larger estimates. The two measurements were only moderately correlated (Pearson r = 0.46; Fig. 1g), and the Bland-Altman plot revealed a mean bias of +0.25 µm with limits of agreement from -0.17 to +1.81 µm (Fig. 1h). These results demonstrate that single plane 2D measurements cannot reliably capture true 3D spine geometry.

### SMH enables reproducible 3D spine morphometry

To obtain orientation-independent measurements, we developed the Spine Morphometry Hub (SMH), a 12-point anatomical landmark framework deployed within the Neuroglancer volumetric viewer on the H01 petavoxel dataset (Fig. 2a-e). The 12 landmarks define the insertion base (P1_BASE), four neck cross-sections (NECK_1A/B, NECK_2A/B), the head–neck junction (P2_UNION), the head centroid (C_HEAD), and five head-surface points (TOP, SIDE1/2, FRONT, BACK). From these coordinates, SMH computes three complementary metric blocks: geometric (analytical from landmarks), voxel-based (from the segmentation mask), and mesh-based (from the LOD-0 triangulated surface; Fig. 2f).

**Fig. 2.**
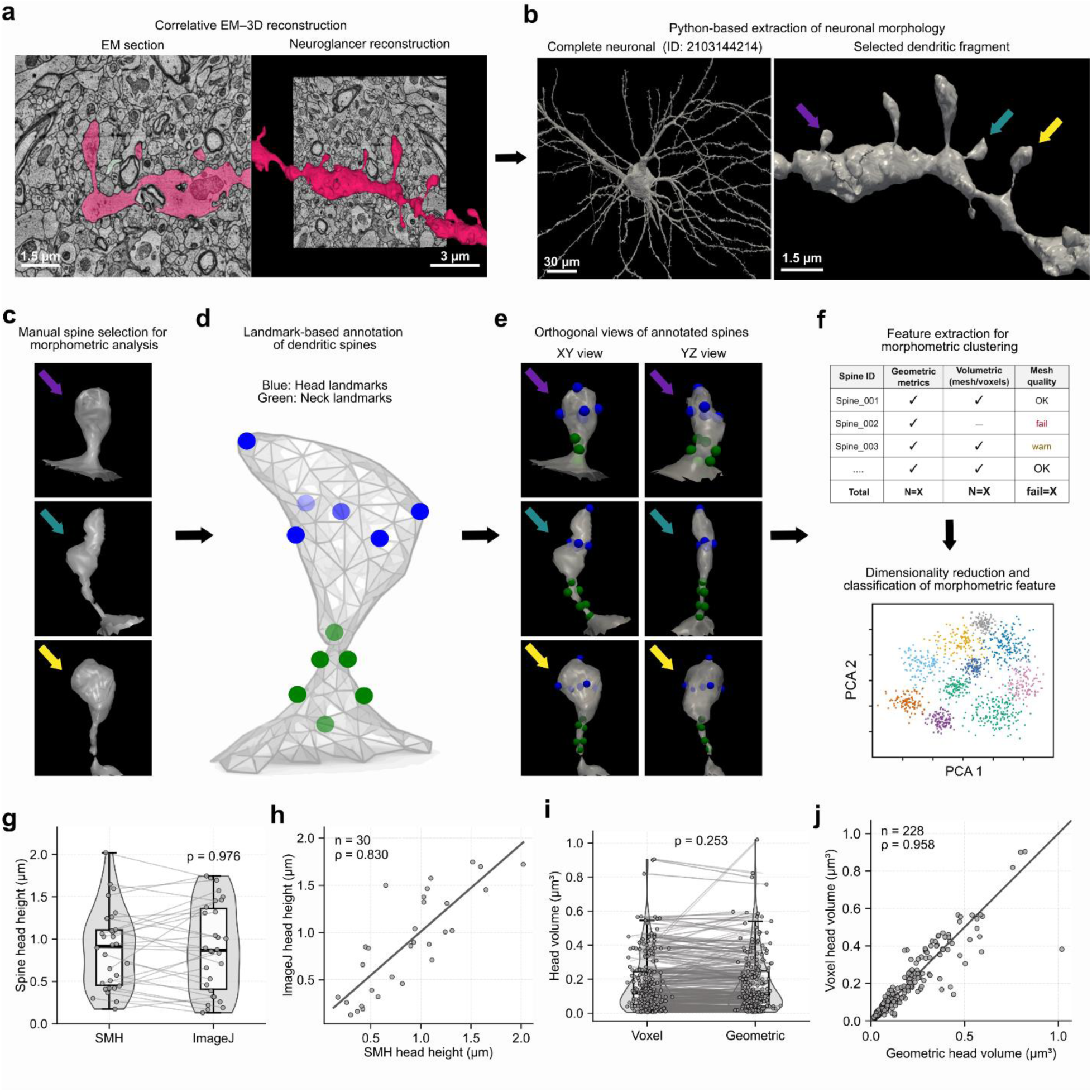
Landmark-based pipeline for orientation-invariant morphometric quantification of dendritic spines. **a**, Correlative workflow: transmission electron micrograph (left; scale bar, 1.3 µm) and corresponding Neuroglancer 3D reconstruction (right; scale bar, 3 µm). **b**, Python-based extraction via CloudVolume. Left: complete neuron reconstruction (scale bar, 30 µm). Right: selected dendritic fragment with representative spines (scale bar, 1.5 µm). **c**, Manual isolation and orientation of individual spines for landmark placement. **d**, The 12-point Spine Morphometry Hub (SMH) framework applied to a single spine: six head landmarks (blue) and six neck/insertion landmarks (green) superimposed on the LOD-0 mesh. **e**, Orthogonal (XY, YZ) views of three annotated spines verifying three-dimensional landmark placement. **f**, Schematic of the feature-extraction and classification workflow, showing the morphometric feature table and PCA scatter plot of the multidimensional morphospace. **g**, Validation of SMH-derived head height against manual ImageJ measurements (n = 30 spines; Wilcoxon signed-rank test, p = 0.976). **h**, Correlation of SMH versus ImageJ head height (n = 30; Pearson r = 0.822, p < 0.001; Spearman ρ = 0.830, p < 0.001). **i**, Comparison of voxel-based and geometric head volume estimates for all vol_viable spines (n = 228; Wilcoxon signed-rank test, p = 0.253). **j**, Correlation between voxel and geometric head volume (n = 228; Pearson r = 0.912, p < 0.001; Spearman ρ = 0.958, p < 0.001; Lin’s CCC = 0.908).

Validation against manual ImageJ measurements on 30 independent spines confirmed SMH reliability. For head height, SMH and ImageJ agreed closely (Wilcoxon signed-rank test, p = 0.976; Fig. 2g), with strong correlation (Pearson r = 0.822, Spearman ρ = 0.830; both p < 0.001; Fig. 2h). Cross-method comparison of voxel-based and geometric head volume estimates across all n = 228 vol_viable spines confirmed tight agreement (Wilcoxon p = 0.253; Pearson r = 0.912, Spearman ρ = 0.958, Lin’s CCC = 0.908; Fig. 2i-j).

### Spine morphometry shows unimodal right-skewed distributions

We analyzed 228 dendritic spines that passed the vol_viable quality-control filter (Table 1). All distributions were right-skewed (Shapiro–Wilk, p < 0.001). Total spine volume distribution had a mean ± s.d. = 0.208 ± 0.182 µm³, median [IQR] = 0.156 [0.065, 0.281] µm³, and range 0.012–1.000 µm³ (Fig. 3a). Head volume had a mean ± s.d. = 0.172 ± 0.167 µm³, median 0.120 [0.042, 0.245] µm³, and range 0.004–0.904 µm³ (Fig. 3b). Head volume and spine volume were strongly correlated (Spearman ρ = 0.975, p < 0.001; Fig. 3c), confirming that head volume is the dominant determinant of total spine volume.

**Table 1.** Orientation-invariant morphometric summary of dendritic spines from human cortical layer II. Summary statistics (mean ± s.d. and range) for five core morphometric variables quantified across n = 228 dendritic spines (vol_viable = 1) from five layer II pyramidal neurons of adult human temporal cortex (H01 dataset). Voxel-derived volumes were computed from the segmentation mask at native resolution (8 × 8 × 33 nm per voxel).

| Variable | N | Mean ± SD | Range |
| --- | --- | --- | --- |
| Head volume (μm³) | 228 | 0.172 ± 0.167 | 0.004–0.904 |
| Spine volume (μm³) | 228 | 0.208 ± 0.182 | 0.012–1.000 |
| Neck length (μm) | 228 | 0.87 ± 0.54 | 0.18–2.94 |
| Neck diameter (μm) | 228 | 0.24 ± 0.12 | 0.08–0.77 |
| Distance soma (μm) | 228 | 154.68 ± 90.68 | 57.67–393.92 |

**Fig. 3.**
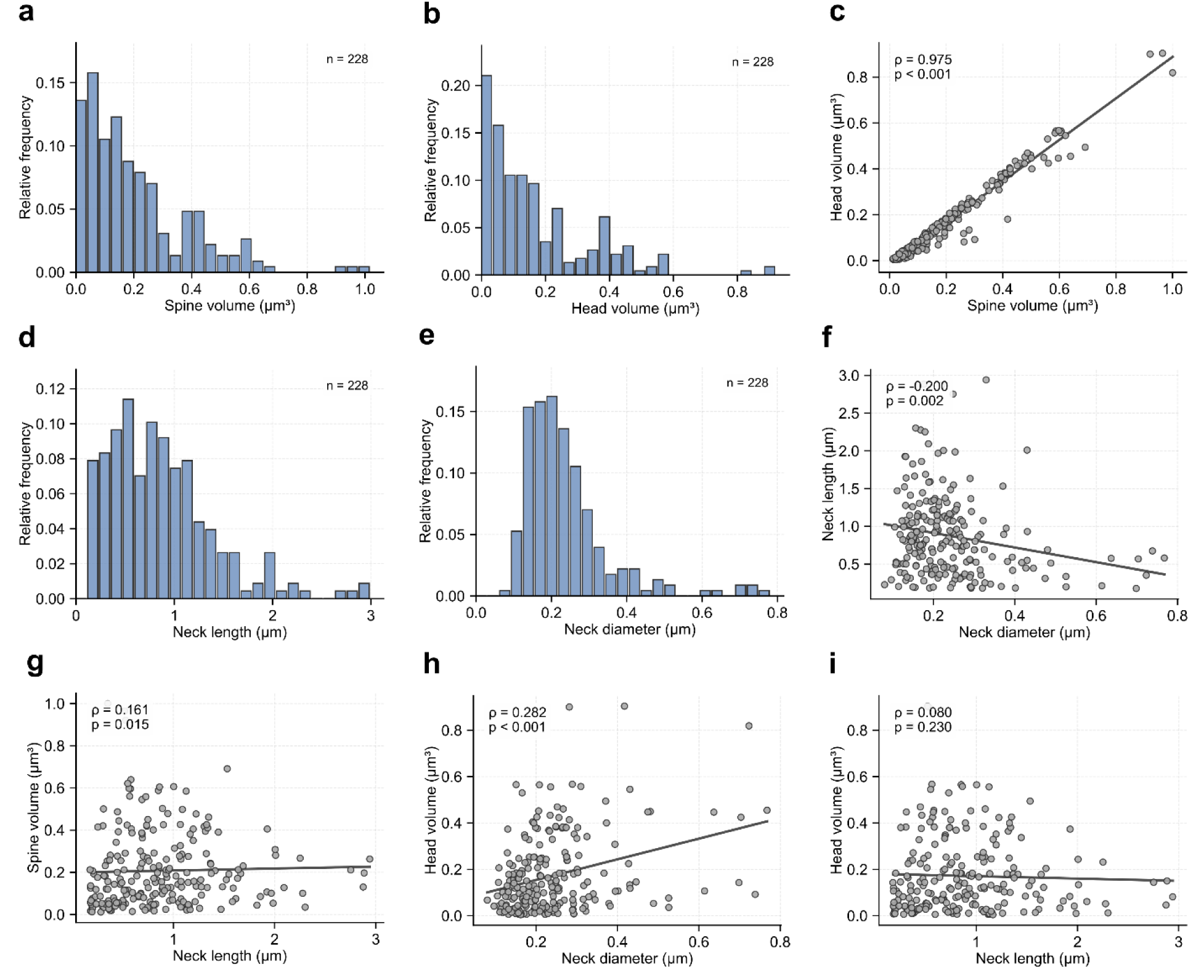
Orientation-invariant morphometric distributions and inter-variable correlations for the 228 dendritic spines. **a-b**, Relative-frequency histograms of spine volume (**a**; mean ± s.d. = 0.208 ± 0.182 µm³, median [IQR] = 0.156 [0.065, 0.281] µm³; Shapiro–Wilk p < 0.001) and head volume (**b**; 0.172 ± 0.167 µm³, median 0.120 [0.042, 0.245] µm³; p < 0.001) for n = 228 spines (vol_viable = 1). **c**, Scatter plot of spine volume versus head volume (Spearman ρ = 0.975, p < 0.001). **d-e**, Histograms of neck length (**d**; 0.874 ± 0.542 µm, median 0.793 [0.471, 1.134] µm) and neck diameter (**e**; 0.240 ± 0.118 µm, median 0.211 [0.163, 0.275] µm). **f**, Neck diameter versus neck length (Spearman ρ = -0.200, p = 0.002). **g**, Neck length versus spine volume (Spearman ρ = 0.161, p = 0.015). **h**, Neck diameter versus head volume (Spearman ρ = 0.282, p < 0.001). **i**, Neck length versus head volume (Spearman ρ = 0.080, p = 0.230, n.s.). Lines in c and f-i are ordinary least-squares fits. All correlations are two-tailed (α = 0.05). n = 228 throughout.

Neck length had a mean ± s.d. = 0.874 ± 0.542 µm, median 0.793 [0.471, 1.134] µm, and range 0.176-2.941 µm (Fig. 3d). Mean neck diameter had a mean ± s.d. = 0.240 ± 0.118 µm, median 0.211 [0.163, 0.275] µm, and range 0.080-0.768 µm (Fig. 3e). Neck diameter and neck length showed a weak inverse relationship (ρ = -0.200, p = 0.002; Fig. 3f). Neck length was weakly but significantly correlated with spine volume (ρ = 0.161, p = 0.015; Fig. 3g), and neck diameter correlated positively with head volume (ρ = 0.282, p < 0.001; Fig. 3h). Neck length and head volume were independent (ρ = 0.080, p = 0.230; Fig. 3i). The full 20-variable Spearman matrix (Fig. 4) confirmed that head-size variables cluster together (ρ ≥ 0.70) while neck geometry varies independently.

**Fig. 4.**
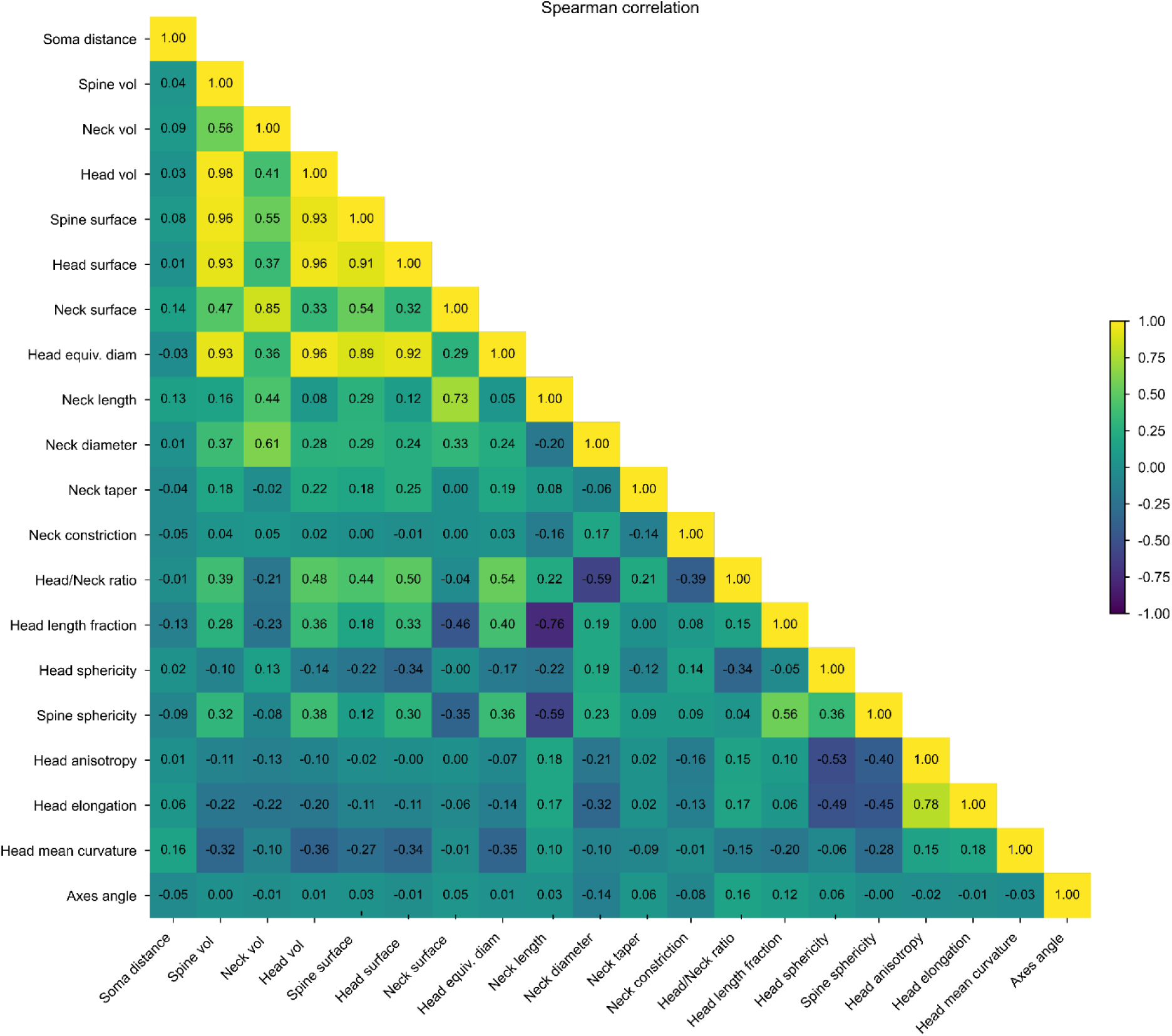
Spearman rank-correlation matrix for 20 orientation-invariant morphometric variables. Lower-triangular Spearman correlation matrix computed for n = 228 spines (vol_viable = 1). Strong positive correlations (ρ ≥ 0.70): head volume vs. spine volume (ρ = 0.975), spine surface vs. spine volume (ρ = 0.960), head surface vs. head volume (ρ = 0.958), head equivalent diameter vs. head volume (ρ = 0.958). Notable negative correlations: head length fraction vs. neck length (ρ = -0.761). All tests were two-tailed (α = 0.05).

A summary of spine morphometry analysis by dendritic compartment is shown in Table 2. Several variables related to the neck, head, spine, geometric characteristics, and location were quantified with our SMH software.

**Table 2.** Orientation-invariant morphometric comparison by dendritic compartment. Summary statistics (mean ± s.d.) for core morphometric variables separated by dendritic compartment (apical, n = 115; basal, n = 113). Comparisons by LMM with neuron_id random intercept; p-values corrected by Benjamini-Hochberg FDR.

| Variable | Mean ± SD<br>All, n=228 | Range | Mean ± SD<br>Apical, n=115 | Range | Mean ± SD<br>Basal, n=113 | Range |
| --- | --- | --- | --- | --- | --- | --- |
| **Neck morphology** |  |  |  |  |  |  |
| Neck length ($\mu\text{m}$) | 0.87 ± 0.54 | 0.18–2.94 | 0.92 ± 0.55 | 0.18–2.88 | 0.83 ± 0.53 | 0.18–2.94 |
| Neck diameter ($\mu\text{m}$) | 0.24 ± 0.12 | 0.08–0.77 | 0.23 ± 0.13 | 0.08–0.77 | 0.24 ± 0.11 | 0.10–0.72 |
| Neck taper ratio | 1.114 ± 0.422 | 0.225–2.918 | 1.106 ± 0.431 | 0.266–2.918 | 1.123 ± 0.413 | 0.225–2.373 |
| Neck constriction index | 0.851 ± 0.118 | 0.367–1.000 | 0.844 ± 0.112 | 0.420–0.999 | 0.857 ± 0.123 | 0.367–1.000 |
| **Head morphology** |  |  |  |  |  |  |
| Head volume ($\mu\text{m}^3$) | 0.172 ± 0.167 | 0.004–0.904 | 0.160 ± 0.163 | 0.004–0.900 | 0.185 ± 0.172 | 0.006–0.904 |
| Head surface area ($\mu\text{m}^2$) | 1.781 ± 1.370 | 0.149–8.043 | 1.627 ± 1.241 | 0.149–6.578 | 1.938 ± 1.479 | 0.153–8.043 |
| Head equiv. diameter ($\mu\text{m}$) | 0.620 ± 0.222 | 0.221–1.249 | 0.600 ± 0.221 | 0.226–1.151 | 0.640 ± 0.222 | 0.221–1.249 |
| Head anisotropy | 0.839 ± 0.103 | 0.276–0.987 | 0.854 ± 0.091 | 0.543–0.987 | 0.823 ± 0.111 | 0.276–0.981 |
| Head sphericity | 0.795 ± 0.134 | 0.099–1.000 | 0.800 ± 0.129 | 0.099–1.000 | 0.789 ± 0.139 | 0.126–1.000 |
| Head aspect ratio (c/a) | 2.759 ± 1.565 | 0.371–13.127 | 2.964 ± 1.754 | 0.526–13.127 | 2.550 ± 1.322 | 0.371–7.118 |
| **Spine overall** |  |  |  |  |  |  |
| Spine volume ($\mu\text{m}^3$) | 0.208 ± 0.182 | 0.012–1.000 | 0.192 ± 0.174 | 0.019–0.921 | 0.223 ± 0.189 | 0.012–1.000 |
| Spine surface area ($\mu\text{m}^2$) | 2.295 ± 1.298 | 0.252–6.240 | 2.198 ± 1.248 | 0.453–6.218 | 2.394 ± 1.345 | 0.252–6.240 |
| Spine total length ($\mu\text{m}$) | 1.81 ± 0.70 | 0.46–4.26 | 1.86 ± 0.70 | 0.60–3.70 | 1.76 ± 0.70 | 0.46–4.26 |
| Spine sphericity | 0.617 ± 0.113 | 0.300–1.000 | 0.604 ± 0.125 | 0.300–1.000 | 0.630 ± 0.099 | 0.358–1.000 |
| Axes angle ($^\circ$) | 29.0 ± 17.9 | 2.2–107.4 | 26.9 ± 18.0 | 2.4–107.4 | 31.1 ± 17.6 | 2.2–87.4 |
| Head/neck diam. ratio | 3.58 ± 1.78 | 0.72–9.28 | 3.58 ± 1.83 | 0.72–9.28 | 3.58 ± 1.74 | 0.78–8.69 |
| **Location** |  |  |  |  |  |  |
| Distance soma ($\mu\text{m}$) | 154.68 ± 90.68 | 57.67–393.92 | 188.85 ± 113.38 | 57.67–393.92 | 119.90 ± 34.09 | 75.86–216.29 |

### Apical and basal spines are morphometrically equivalent

Six core morphometric variables were compared between apical (n = 115) and basal (n = 113) spines using linear mixed-effects models (LMMs) with neuron identity as a random intercept (5 neurons), followed by Benjamini-Hochberg FDR correction across 23 variables (Fig. 5a-f). No core variable differed significantly: head volume (q = 0.312, Cohen’s d = -0.15), spine volume (q = 0.289, d = -0.17), neck diameter (q = 0.579, d = - 0.09), neck length (q = 0.339, d = 0.16), neck taper ratio (q = 0.798, d = -0.04) and head/neck diameter ratio (q = 0.998, d < 0.01). All effect sizes were negligible (|d| < 0.20).

**Fig. 5.**
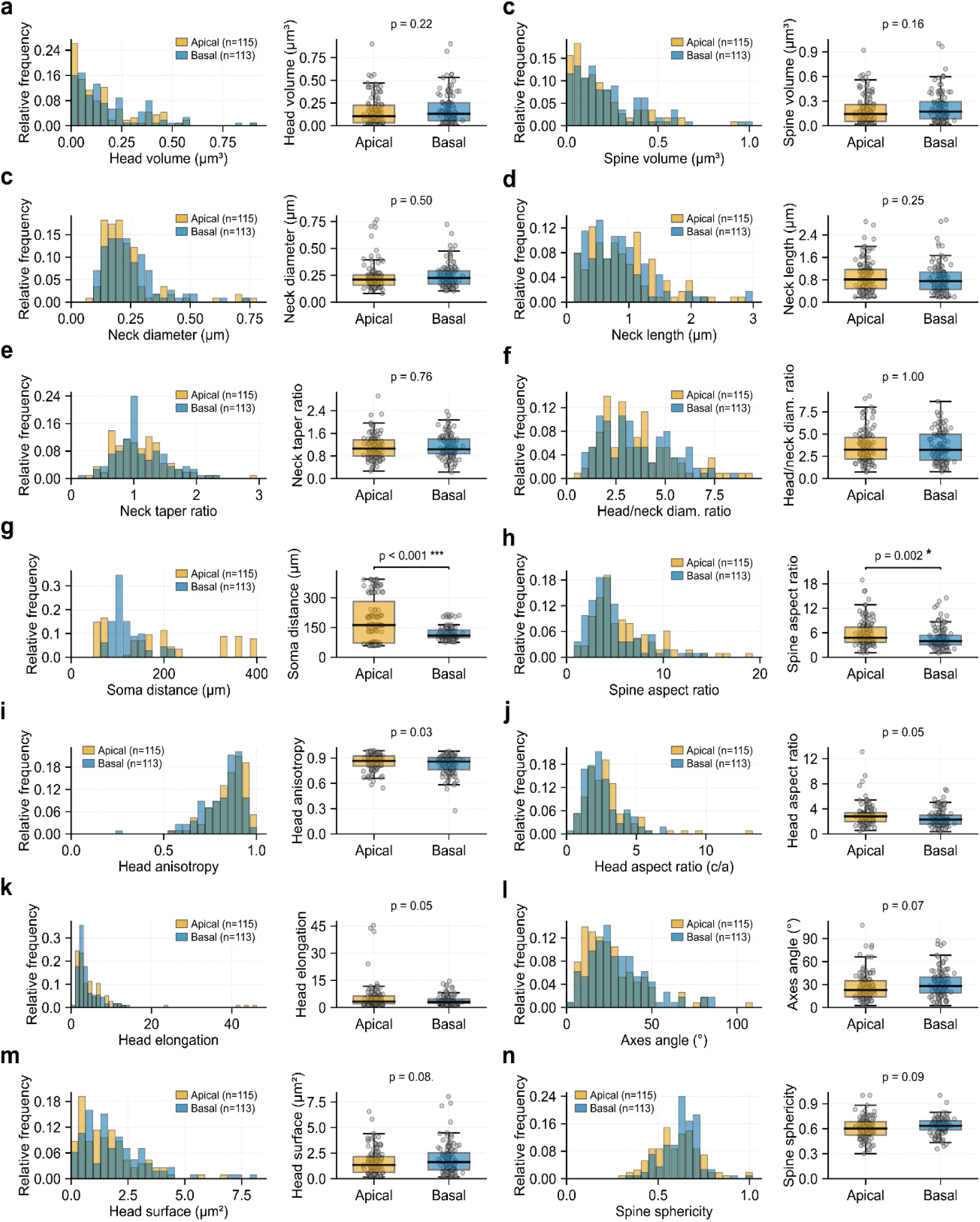
Orientation-invariant morphometric comparison between apical and basal dendritic spines. **a-f**, Boxplots comparing six core variables between apical (orange, n = 115) and basal (blue, n = 113) spines: head volume (**a**), spine volume (**b**), neck diameter (**c**), neck length (d), neck taper ratio (e), and head/neck diameter ratio (**f**). Statistical comparisons were performed using linear mixed-effects models (LMMs) with neuron identity as a random intercept (5 neurons, 228 spines), followed by Benjamini-Hochberg FDR correction across 23 variables. None of the six core variables differed significantly between compartments after FDR correction: head volume (p = 0.312, d = -0.15), spine volume (p = 0.289, d = -0.17), neck diameter (p = 0.579, d = -0.09), neck length (p = 0.339, d = 0.16), neck taper ratio (p = 0.798, d = -0.04), head/neck ratio (p = 0.998, d < 0.01). Two variables reached significance: soma distance (p < 0.001, d = 0.82, large) and spine aspect ratio (p = 0.020, d = 0.42, small). All effect sizes for the six core variables were negligible (|d| < 0.20). **g-n**, Histograms and boxplots for eight additional variables. Soma distance (**g**) was greater in apical spines, reflecting the longer reach of apical dendrites (p < 0.001, d = 0.82, large). Spine aspect ratio (**h**) was higher apically (p = 0.002, d = 0.42, small). The remaining variables showed no significant compartmental difference after FDR correction: (**i**) head anisotropy (p = 0.03), (**j**) head aspect ratio (p = 0.05), (**k**) head elongation (p = 0.05), (**l**) axes angle (p = 0.07), (**m**) head surface area (p = 0.08), and (**n**) spine sphericity (p = 0.09). Apical, orange; basal, blue; n = 115 and n = 113, respectively, throughout.

Two variables reached significance after FDR correction: soma distance (q < 0.001, d = 0.82, large effect; apical 188.9 ± 113.4 µm vs basal 119.9 ± 34.1 µm), reflecting the greater length of apical dendrites, and spine aspect ratio (q = 0.020, d = 0.42, small effect; apical 5.93 ± 3.30 vs basal 4.67 ± 2.66). The ICC for soma distance was 0.563, indicating substantial between-neuron variance; all other ICCs were below 0.06. These results confirm that spine size and neck geometry are statistically indistinguishable between dendritic compartments.

### Spine morphology forms a continuous multidimensional space

PCA was applied to seven morphometrically independent variables (neck length, neck diameter, neck taper ratio, head volume, head sphericity, head/neck diameter ratio, and axes angle; RobustScaler; n = 228). Six components were required to explain ≥90% of variance (cumulative variance at PC6 = 97.7%; Fig. 6a). PC1 accounted for 29.9%, PC2 for 20.7%, and PC3 for 15.5% (Fig. 6b). The effective dimensionality was D_eff = 5.25 out of 7, confirming that morphological variation is distributed across multiple axes.

**Fig. 6.**
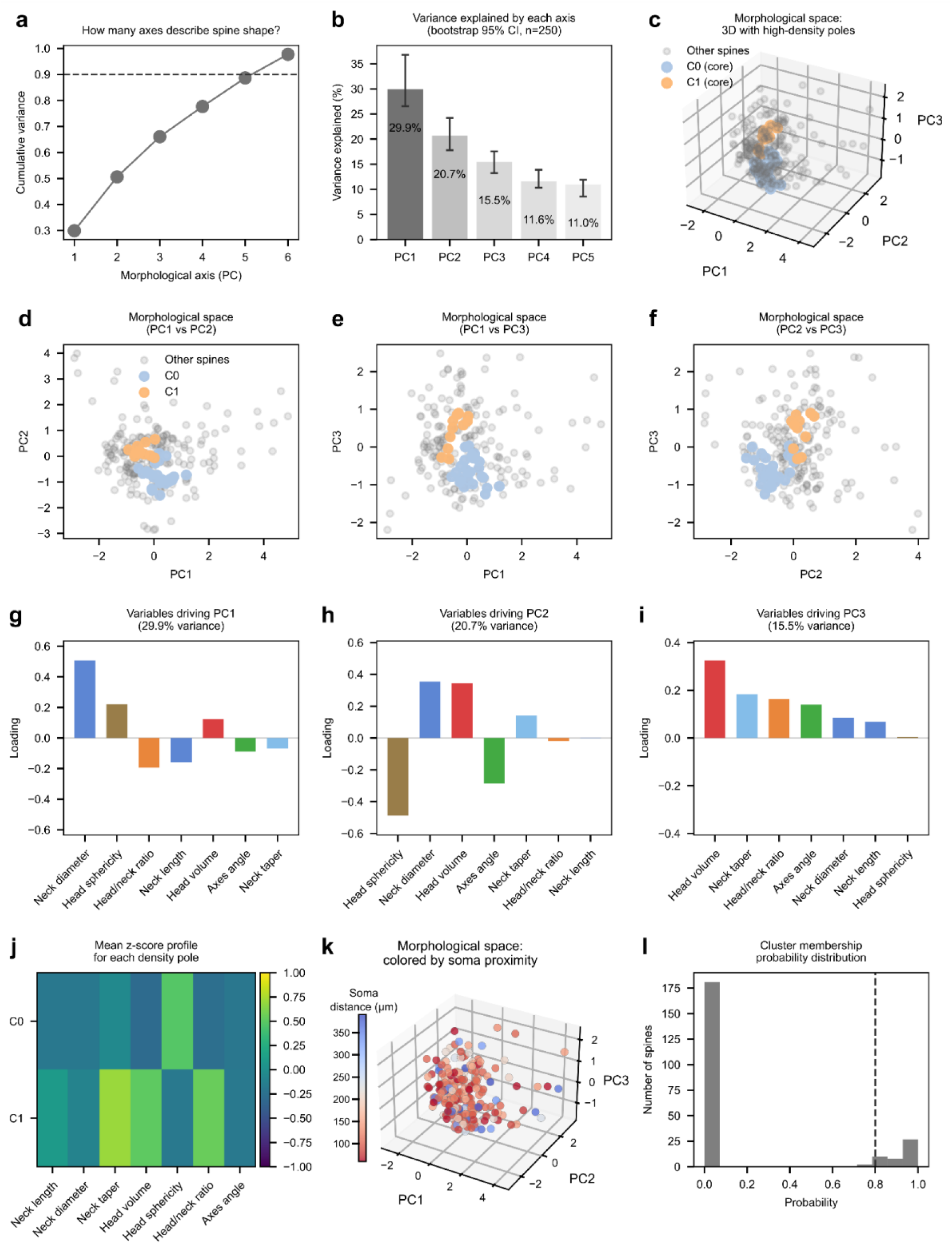
A continuous orientation-invariant morphological space reveals the absence of discrete spine subtypes. **a**, Cumulative variance explained by principal components derived from seven morphometric variables (neck length, neck diameter, neck taper ratio, head volume, head sphericity, head/neck diameter ratio, axes angle; RobustScaler; n = 228). Six components capture 97.7% of the variance. **b**, Variance explained by PC1-PC3 with bootstrap 95% CIs (250 resamples). **c**, Three-dimensional PC1-PC3 morphospace colored by PC1 score gradient. HDBSCAN-assigned spines at full opacity; unassigned spines (n = 181, 79.4%) at reduced opacity. **d-f**, Two-dimensional projections with KDE contours: PC1-PC2 (**d**), PC1-PC3 (**e**), PC2-PC3 (**f**). **g-i**, Loading bar charts for PC1 (**g**), PC2 (**h**), and PC3 (**i**). **j**, Mean standardized z-scores at PC1 extremes (Q10 vs Q90). **k**, KDE density landscape in PC1-PC2 space. **l**, Score distributions along PC1-PC3. HDBSCAN (min_cluster_size = 7, min_samples = 3, eom) identified two clusters: C0 (n = 36) and C1 (n = 11), with 181 spines (79.4%) unassigned, consistent with a morphological continuum rather than discrete subtypes.

PC1 loadings were dominated by head volume, head/neck diameter ratio, and head sphericity, representing a head-size axis; PC2 captured independent neck-geometry variation (neck length, neck taper); PC3 reflected the three-dimensional axis angle (Fig. 6g-i). Comparison of mean standardized z-scores at the extremes of the morphological continuum (Q10 vs Q90 of PC1 scores) confirmed that PC1 separates small thin protrusions from large mushroom-like spines (Fig. 6j).

The PC1-PC3 morphospace showed spines distributed as a continuous, elongated cloud with no visible gaps or discrete groupings (Fig. 6c-f). KDE density landscapes confirmed a smooth, unimodal distribution (Fig. 6k). Score distributions along all three axes were unimodal (Fig. 6l).

HDBSCAN (min_cluster_size = 7, min_samples = 3, eom) assigned 181 spines (79.4%) to noise and identified two high-density regions: C0 (n = 36) and C1 (n = 11; silhouette coefficient = 0.285; Fig. 6c). The high noise fraction indicates that many spines do not belong to any discrete cluster but instead occupy a broad, low-density continuum. Examination of mean z-score profiles revealed that C0 comprises small-headed spines with short, thin necks (low PC1 and PC2 scores), whereas C1 comprises large-headed spines with long, wide necks (high PC1, moderate PC2). These regions represent the most frequently visited morphological configurations along a continuous gradient that reflects the population’s distributional extremes, rather than biologically distinct spine subtypes.

To test whether the high noise rate reflects genuine biological continuity rather than algorithmic sensitivity, we performed a Monte Carlo simulation (1,000 iterations). Synthetic datasets were drawn from a Gaussian mixture model with k = 2 components (BIC-selected) and within-cluster spread reduced to 30% of the empirical standard deviation (Fig. 7). Under this null hypothesis of two discrete types, the mean noise rate was 10.3% ± 8.1% (95% CI: 0.9-31.1%), far below the 76.3% observed in the real data (p < 0.001). This confirms that the observed morphological distribution is inconsistent with discrete categories.

**Fig. 7.**
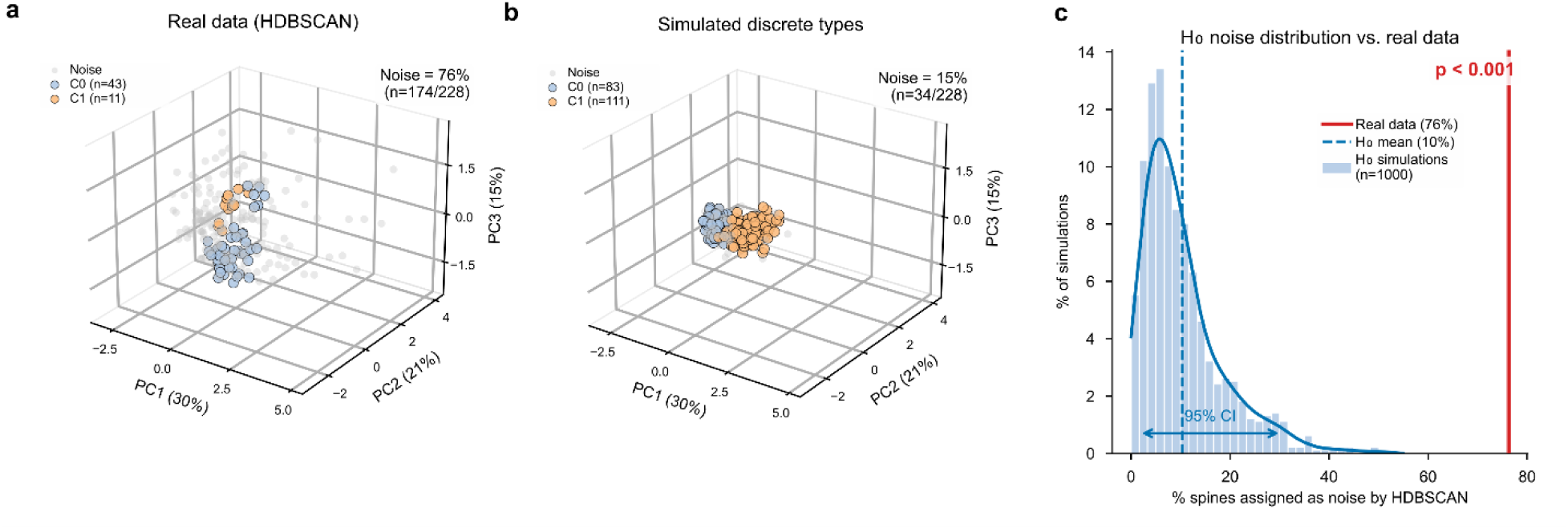
Monte Carlo simulation under the null hypothesis of discrete spine types. **a**, PC1-PC2 projection of n = 228 spines with HDBSCAN assignments. Noise (grey, n = 174, 76.3%); C0 and C1 color-coded. **b**, Representative synthetic dataset under H₀ (GMM with k = 2, BIC-selected; within-cluster SD = 30% of empirical). HDBSCAN recovers discrete groups with minimal noise. **c**, Distribution of noise rates across 1,000 simulations: mean = 10.3% ± 8.1% (95% CI: 0.9–31.1%). The observed 76.3% noise rate (red line) exceeds all simulated values (p < 0.001), confirming that the high noise rate reflects genuine morphological continuity.

### Morphological space is independent of the dendritic compartment

To test whether compartment identity is detectable in the multivariate morphospace, apical (n = 115) and basal (n = 113) spines were compared in PC1-PC3 space (Fig. 8a). Both HDBSCAN clusters contained spines from both compartments: C0 (19 apical, 17 basal) and C1 (4 apical, 7 basal; Fig. 8b). A logistic regression classifier trained on PC1-PC3 scores (5-fold stratified cross-validation) yielded AUC = 0.461, at chance level, confirming that compartment cannot be inferred from morphometric features (Fig. 8c).

**Fig. 8.**
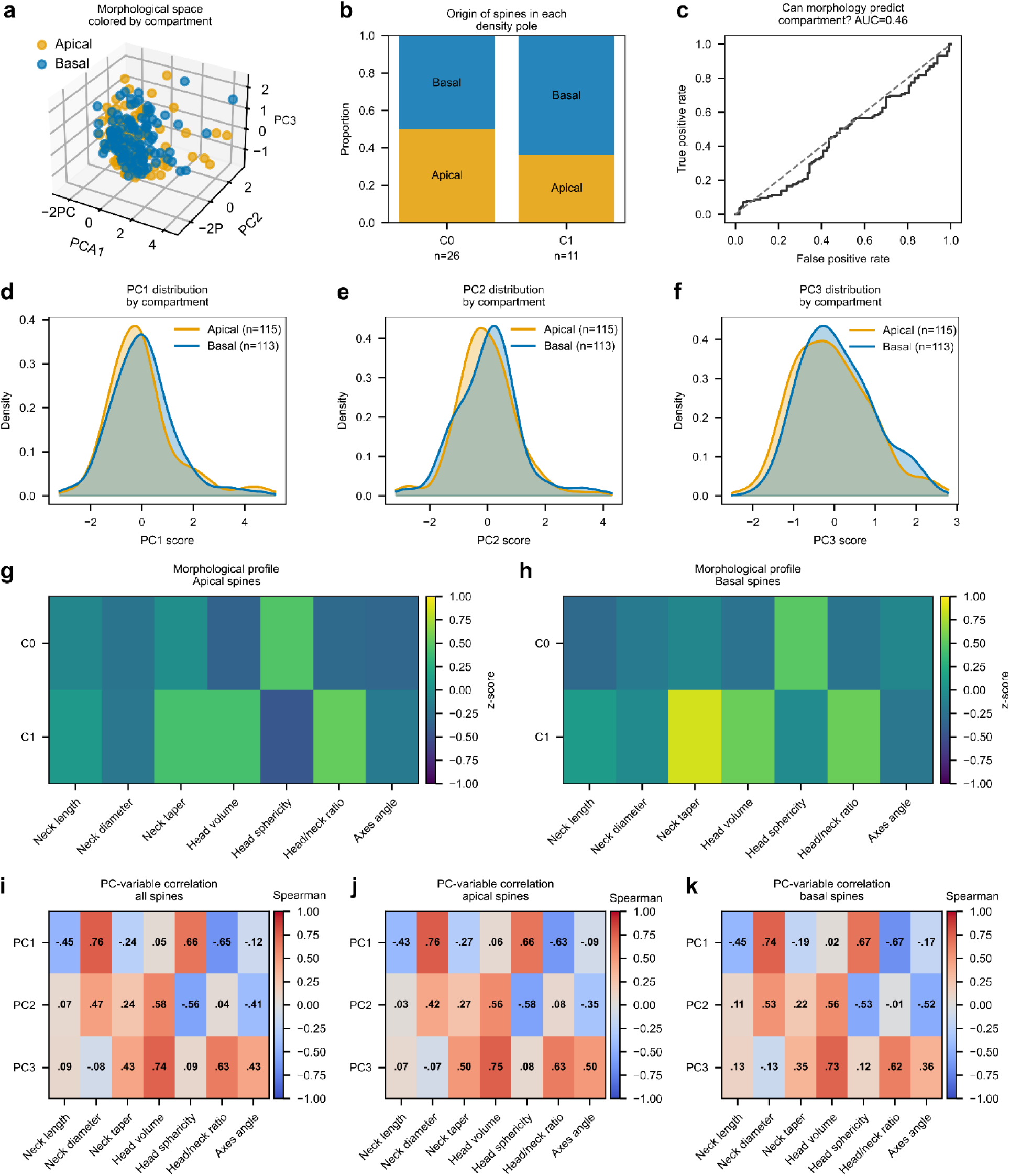
Compartment origin does not determine position in the spine orientation-invariant morphological space. **a**, PC1-PC3 morphospace (n = 228) colored by compartment: apical (orange, n = 115) and basal (blue, n = 113). Both compartments overlap extensively. **b**, Composition of HDBSCAN clusters by compartment. C0 (n = 36: 19 apical, 17 basal) and C1 (n = 11: 4 apical, 7 basal) contain spines from both compartments. c, ROC curve for logistic regression PC1-PC3) discriminating apical from basal spines (5-fold stratified cross-validation). **d-f**, Overlapping PC score histograms for apical and basal spines. **g-h**, Heatmaps of mean z-scores per cluster for apical (**g**) and basal (**h**) subsets. **i-k**, Spearman correlation heatmaps computed globally (**i**) and separately for apical (**j**) and basal (**k**) subsets. The correlation structure is conserved across compartments.

Both compartments overlapped extensively across the entire morphospace. Kolmogorov-Smirnov tests on individual PC scores revealed no significant differences (PC1: D = 0.146, p = 0.151; PC2: D = 0.141, p = 0.183; PC3: D = 0.102, p = 0.536; Fig. 8d-f).

Spearman correlation heatmaps, computed separately for apical and basal subsets, showed a conserved correlation structure (Fig. 8i-k), and mean z-score profiles per cluster were preserved in both compartments (Fig. 8g,h).

## Discussion

The continuous organization of dendritic spine morphology has been reported before in rodent and human cortex alike, but always with residual doubt. If two-dimensional measurements are unreliable, the apparent continuum might itself be a measurement artifact rather than a property of the tissue. A previous study (Manjarrez et al., 2026) sharpened that doubt by showing that categorical assignment of human spines barely survives a change of viewing angle (Cohen’s κ = 0.027). The present work resolves it. By measuring 228 spines in three orthogonal planes with a landmark framework that is invariant to dendritic orientation, we remove the very source of instability and ask whether discrete subtypes re-emerge once the artifact is gone. They do not. The morphometric distributions remain unimodal and right-skewed, density-based clustering assigns the great majority of spines to no cluster at all, and a Monte Carlo test calibrated against a deliberately compact two-type null rejects the discrete model outright. The continuum, in other words, is what remains after the measurement error is removed, not what the error produced. What our analysis contributes beyond confirmation is, therefore, methodological as much as conceptual. We provide a quantification of the orientation-dependent error itself, a multi-block morphometric framework with internal cross-validation, and a formal statistical criterion for distinguishing a continuum from a set of categories.

The quantification of orientation-dependent error addresses a long-standing but underappreciated methodological concern. The bidirectional nature of the error is striking: a 90° rotation of the dendritic axis produced statistically significant increases in apparent height in half of the spines and decreases in the other half, with no net group-level bias. This pattern reflects the geometric reality that a 2D projection captures only the component of the spine vector lying in the imaging plane. The 3.52 µm span of the Bland-Altman limits of agreement for height and the systematic plane-dependent bias for head width indicate that neither Golgi-impregnated material examined by light microscopy nor confocal fluorescence imaging with axial resolution of 600-800 nm can provide reliable individual-spine measurements. This complements the findings of Kashiwagi et al. (2019), who demonstrated that structured illumination microscopy achieves comparable accuracy for horizontally and vertically protruding spines, but that 2D projection cannot. Volume electron microscopy eliminates this limitation by providing near-isotropic nanometer-scale resolution in all three dimensions (Lichtman & Denk, 2011). The implication for existing literature is that published studies relying on 2D categorical classification of spine type in human postmortem tissue incur orientation-induced misclassification rates that cannot be estimated retrospectively.

The Spine Morphometry Hub addresses a gap between the resolution of modern connectomic datasets and the tools available to exploit them for quantitative morphometry. Existing automated pipelines such as DeepD3 (Fernholz et al., 2024), SpineJ (Levet et al., 2020), and the framework of Bernal-Garcia et al. (2025) operate on fluorescence microscopy data and are constrained by diffraction-limited resolution and the assumptions inherent in template-based shape fitting. Luengo-Sanchez et al. (2018) applied clustering approaches to over 7,000 human pyramidal cell spines reconstructed from confocal microscopy, confirming continuous morphological variation, but their measurements remained subject to the optical resolution limit and did not separate geometric, voxel, and mesh metric blocks. SMH operates directly on volume EM data at the native voxel resolution of the connectome, extracts three independent and complementary metric blocks from the same physical object, and implements explicit quality-control criteria that select spines based on volumetric concordance across methods rather than categorical shape assumptions.

The close agreement between voxel-based and geometric head volume estimates across the full sample validates this multi-block approach and provides internal consistency checks absent from single-method pipelines. The availability of the H01 petavoxel dataset (Shapson-Coe et al., 2024), the first nanometer-resolution reconstruction of a cubic millimeter of human neocortex, was essential for this work: it provides the segmentation masks, surface meshes, and native-resolution imagery needed to extract three independent metric blocks from the same physical spine, a capability that no other publicly available human dataset currently offers.

The continuous distribution of spine morphology in 228 human cortical spines converges with and extends ultrastructural evidence from rodent tissue. Arellano et al. (2007) examined 144 completely reconstructed spines from mouse visual cortex layer II/III and concluded that no clear subgrouping could be detected, with all distributions single-peaked and right-skewed. Ofer et al. (2021) demonstrated a morphological continuum in spine neck geometry, and Ofer et al. (2022) provided direct evidence for a continuum in human cortical spines using three-dimensional reconstructions. Kashiwagi et al. (2019) reached the same conclusion using PCA on five shape descriptors in cultured hippocampal neurons.

Parallel ultrastructural work from the DeFelipe laboratory has provided complementary population-level evidence in human tissue: three-dimensional analyses of synaptic organization in temporal and trans entorhinal cortex (Dominguez-Alvaro et al., 2021; Montero-Crespo et al., 2020, 2021; Cano-Astorga et al., 2021, 2024) have characterized spine and synapse geometry without requiring discrete categorical boundaries, and variation in pyramidal cell morphology across cortical regions further supports a continuous gradient (Benavides-Piccione et al., 2021).

Our analysis advances this body of work in three respects: it is performed on adult human tissue at nanometer resolution; it employs seven functionally independent variables that capture distinct structural dimensions; and it applies formal density-based topology testing with Monte Carlo calibration rather than visual inspection or k-means partitioning. The convergence of an effective dimensionality of 5.25 out of 7, unimodal score distributions along all retained principal components, a silhouette coefficient of 0.285 for the two identified high-density regions, and the failure of a supervised classifier to discriminate the compartment of origin collectively make it unlikely that the continuous distribution is an artifact of any single analytical choice.

The continuous morphological distribution is the expected outcome of the biology of activity-dependent structural plasticity. Long-term potentiation is accompanied by graded, stimulus-proportional head enlargement rather than a discrete category switch (Matsuzaki et al., 2004; Holtmaat & Svoboda, 2009; Nicoll & Schulman, 2023; Jain et al., 2024). Long-term depression produces the reverse: gradual head shrinkage and neck elongation along the same continuous dimensions (Kasai et al., 2010; Steffens et al., 2021).

If the mechanism of morphological change is intrinsically continuous, the resulting population-level distribution must also be continuous. Discrete categories would require deep attractor states with high energy barriers between them, yet the dominant picture from population-level ultrastructure in adult neocortex is of unimodal distributions with broad variance (Nimchinsky et al., 2002; Kasai et al., 2010). Within this framework, the two high-density regions identified by HDBSCAN represent the morphological configurations most frequently visited under the current activity regime, not discrete biological subtypes. The classical tripartite classification can be reinterpreted as a coarse discretization of a smooth gradient along the first principal axis.

The near-zero correlation between neck length and head volume and the weak but significant inverse relationship between neck diameter and neck length are consistent with the emerging molecular picture in which head growth and neck remodeling are controlled by partially distinct signaling cascades. Head volume is regulated predominantly by AMPA receptor insertion and by Rac1-driven F-actin polymerization (Matsuzaki et al., 2001; Hotulainen & Hoogenraad, 2010). Neck geometry is subject to independent modulation by myosin IIb contractility and cofilin-mediated actin severing (Tonnesen et al., 2014; Araya et al., 2014). Koch and Zador (1993) showed that neck resistance determines the degree of electrical and biochemical compartmentalization.

Because neck geometry and head size vary independently in our population, spines with identical synaptic strength can differ markedly in their degree of compartmentalization, and spines with equivalent compartmentalization can differ in synaptic strength. This geometric diversity in the compartmentalization–strength plane may provide a substrate for context-dependent plasticity rules.

The absence of morphometric differences between apical and basal spines after correction for pseudo replication is notable and consistent with previous studies (Arellano et al., 2007), given the well-documented differences in input statistics, dendritic integration properties, and plasticity rules between these compartments (Spruston, 2008; Larkum et al., 1999). In our data, the only variables that distinguished compartments were soma distance, which reflects dendritic length rather than spine geometry, and spine aspect ratio, a secondary shape descriptor with a small effect size. This result is also consistent with Arellano et al. (2007), who found no significant compartmental differences in mouse neocortex.

Benavides-Piccione et al. (2013, 2024) reported similar equivalence in three-dimensional reconstructions of human cingulate and temporal cortex, and systematic cross-regional comparisons of pyramidal cell morphology and spine dimensions (Benavides-Piccione et al., 2021, 2025) suggest that morphometric equivalence between apical and basal spines is a conserved feature of human pyramidal neurons. The most parsimonious interpretation is that individual spine morphology is determined primarily by the history of local synaptic activity rather than by compartment-specific transcriptional programs (Harvey & Svoboda, 2007).

The present work extends the findings of Yuste, Arellano, and colleagues in several important respects. Arellano et al. (2007) established the morphological continuum in the mouse cortex using serial-section electron microscopy with a sample of 144 spines and five morphometric variables. Ofer et al. (2021, 2022), working within the same group, confirmed this continuum for neck geometry and extended it to human cortical spines using three-dimensional confocal reconstructions. Yuste (2011) proposed that spines serve as distributed circuit elements whose individual geometry reflects local synaptic history. Yuste (2013) formalized the concept of electrical compartmentalization in spines, and Cornejo et al. (2022) demonstrated that spines function as independent voltage compartments in vivo.

Our study advances beyond these contributions in five ways. First, we provide the first systematic quantification of orientation-dependent measurement error in human tissue, demonstrating that the 2D projection methods used in the majority of the existing spine morphometry literature produce individual-spine errors exceeding six times the mean head diameter; this finding has not been reported previously and calls into question the reliability of categorical morphometry based on 2D measurements.

Second, whereas previous studies relied on independent geometric or voxel-based measurements, SMH integrates three complementary metric blocks from the same physical object with explicit cross-method concordance filtering, providing internal validation absent from prior work.

Third, we apply linear mixed-effects models with neuron identity as a random intercept and false discovery rate correction across all variables, addressing the pseudo replication inherent in sampling multiple spines from the same neurons, a statistical concern not accounted for in prior studies that used Mann–Whitney U tests without clustering correction.

Fourth, the Monte Carlo null hypothesis test with BIC-selected Gaussian mixture components provides a formal statistical framework for evaluating the HDBSCAN noise rate against a calibrated null distribution, replacing the qualitative arguments for continuity offered in earlier work.

Fifth, the combination of SMH with the H01 petavoxel dataset enables morphometric analysis at the native voxel resolution of 8 × 8 × 33 nm, an order of magnitude finer than diffraction-limited optical methods, applied for the first time to a systematic population of human cortical spines.

### Limitations

However, our study has several limitations. Although the number of spines is high, the analysis is restricted to five neurons from a single cortical region. Whether the continuous morphological space and compartmental equivalence generalize across cortical areas, ages, or disease states requires investigation using larger, more diverse samples.

The anisotropic voxel resolution of the H01 dataset (33 nm in Z versus 8 nm in X and Y) may introduce minor systematic error in volume estimates for spines aligned with the Z-axis, although the close agreement between voxel-derived and geometric volumes across the sample suggests that this effect is small. The landmark protocol relies on a single annotator; formal multi-annotator reliability studies are needed. Moreover, volume electron microscopy captures a single time point in the dynamic life of structurally plastic spines (Tonnesen et al., 2014; Bhatt et al., 2009), and the morphospace we observe may partly reflect the distribution of recent activity states.

Chemical fixation with glutaraldehyde and osmium tetroxide may produce differential shrinkage at the nanometer scale, and the magnitude of these effects in the H01 preparation has not been validated against cryo-electron microscopy.

### Future directions

Despite these limitations, establishing a quantitative, orientation-invariant baseline for human cortical spine morphometry has direct implications for neuropathological research. Spine pathology is among the most consistently reported anatomical correlates of cognitive impairment in Alzheimer’s disease, schizophrenia, and neurodevelopmental disorders (Terry et al., 1991; Glantz & Lewis, 2000; Glausier & Lewis, 2013; Penzes et al., 2011). Regional specificity of spine loss has been demonstrated across cortical layers and areas in schizophrenia (Fish et al., 2025), and altered synaptic ultrastructure has been documented in animal models of autism spectrum disorder (Jacot-Descombes et al., 2020). Walker et al. (2024) demonstrated that spine head diameter in human postmortem tissue predicts episodic memory performance. The SMH framework enables quantitative assessment of whether disease processes displace spines toward specific regions of the continuous morphospace.

Because apical and basal spines occupy the same morphological territory in unaffected tissue, any compartment-specific pathological change would constitute a positive finding. The transition from categorical to continuous morphometric endpoints would improve statistical power and enable the detection of subtle drug effects below the threshold for categorical reclassification. In summary, this study provides the first validated orientation-invariant morphometric framework for human cortical dendritic spines at nanometer resolution and demonstrates that morphological diversity is continuous, broadly distributed, and independent of dendritic compartment. These findings reframe the classical discrete classification as a coarse approximation of a smooth morphological gradient and establish the infrastructure for comparative studies in health and disease.

Finally, electron microscopy datasets cannot be combined with functional recordings from the same preparation; correlating morphological position with synaptic physiology will require future correlative approaches or large-scale electron microscopy of functionally characterized circuits (Turner et al., 2022).

### Comparative summary of our results with previous studies to validate our orientation-invariant morphometry framework

In summary, we have three principal findings: (i) we confirm that two-dimensional projections introduce biologically significant orientation-dependent measurement error, as demonstrated by Manjarrez et al., 2026; (ii) we confirm that human cortical dendritic spines occupy a continuous, broadly distributed morphological space in which density-based clustering assigns 79% of spines to noise under formal Monte Carlo calibration (as demonstrated in previous electron microscopy and confocal studies by Arellano et al., 2007 and Ofer et al., 2022); and (iii) we confirm that apical and basal spines are morphometrically indistinguishable when pseudo replication is controlled using linear mixed-effects models with false discovery rate correction, as demonstrated by Ofer et al., 2022. These findings validate our orientation-invariant morphometry framework for assessing pathological spine remodeling in neurological and psychiatric disorders.

### Conclusion

For more than a century, dendritic spines have been sorted into types because that is how the eye sorts them under a microscope. The categories were never in the tissue. They came from the way we looked at it. When we control the angle of view and measure each spine in three dimensions, the borders between types disappear, and we are left with a continuous range of shapes that a neuron can move through, one spine at a time. This changes what it means to study spine pathology. A diseased cortex does not hold more spines of one class and fewer of another, since those classes were never real to begin with. It holds a population that has moved to a different visualization, with continuous orientation-invariant morphometry, as the challenge confronting the field.

## Author contributions

Conceptualization (EM); Data curation (MAZU and EM); Formal analysis (MAZU and EM); Funding acquisition (EM); Investigation (MAZU and EM); Methodology (MAZU and EM); Software (MAZU and EM); Validation (EM); Writing - original draft (MAZU); Writing - review and editing (EM).

## Ethical Approval Statement

This manuscript does not require an ethical approval statement, as it was performed on a publicly available database.

## Data Availability Statement

The H01 dataset is publicly available at https://h01-release.storage.googleapis.com. The complete morphometric dataset (metrics CSV with all 228 annotated spines and viability flags) and the 12-point landmark coordinates will be deposited at [URL/DOI https://github.com/ to be provided upon acceptance].

## Acknowledgements

We gratefully acknowledge the H01 consortium (J. W. Lichtman, V. Jain, A. Shapson-Coe, M. Januszewski, and colleagues) for generating and publicly releasing the petavoxel human cortex dataset (Shapson-Coe et al., 2024), without which the present analyses would not have been possible. We also thank the developers of Neuroglancer, CloudVolume, and the open-source Python scientific computing ecosystem. MAZU acknowledges Postdoc support from SECIHTI. EM acknowledges support from the Marcos Moshinsky Foundation and VIEP-BUAP.

## References

Alvarez, V. A., & Sabatini, B. L. (2007). Anatomical and physiological plasticity of dendritic spines. Annual review of neuroscience, 30, 79–97. 10.1146/annurev.neuro.30.051606.094222

Araya, R., Vogels, T. P., & Yuste, R. (2014). Activity-dependent dendritic spine neck changes are correlated with synaptic strength. Proceedings of the National Academy of Sciences of the United States of America, 111(28), E2895–E2904. 10.1073/pnas.1321869111

Arellano, J. I., Benavides-Piccione, R., Defelipe, J., & Yuste, R. (2007). Ultrastructure of dendritic spines: correlation between synaptic and spine morphologies. Frontiers in neuroscience, 1(1), 131–143. 10.3389/neuro.01.1.1.010.2007

Benavides-Piccione, R., Blazquez-Llorca, L., Kastanauskaite, A., Fernaud-Espinosa, I., Tapia-González, S., & DeFelipe, J. (2024). Key morphological features of human pyramidal neurons. Cerebral cortex (New York, N.Y. : 1991), 34(5), bhae180. 10.1093/cercor/bhae180

Benavides-Piccione, R., Fernaud-Espinosa, I., Kastanauskaite, A., & DeFelipe, J. (2025). Principles for Dendritic Spine Size and Density in Human and Mouse Cortical Pyramidal Neurons. The Journal of comparative neurology, 533(6), e70060. 10.1002/cne.70060

Benavides-Piccione, R., Fernaud-Espinosa, I., Robles, V., Yuste, R., & DeFelipe, J. (2013). Age-based comparison of human dendritic spine structure using complete three-dimensional reconstructions. Cerebral cortex (New York, N.Y. : 1991), 23(8), 1798–1810. 10.1093/cercor/bhs154

Benavides-Piccione, R., Rojo, C., Kastanauskaite, A., & DeFelipe, J. (2021). Variation in Pyramidal Cell Morphology Across the Human Anterior Temporal Lobe. Cerebral cortex (New York, N.Y. : 1991), 31(8), 3592–3609. 10.1093/cercor/bhab034

Bernal-Garcia, S., Schlotter, A. P., Pereira, D. B., Recupero, A. J., Polleux, F., & Hammond, L. A. (2025). A deep learning pipeline for accurate and automated restoration, segmentation, and quantification of dendritic spines. Cell reports methods, 5(10), 101179. 10.1016/j.crmeth.2025.101179

Bhatt, D. H., Zhang, S., & Gan, W. B. (2009). Dendritic spine dynamics. Annual review of physiology, 71, 261–282. 10.1146/annurev.physiol.010908.163140

Bock, D. D., Lee, W. C., Kerlin, A. M., Andermann, M. L., Hood, G., Wetzel, A. W., Yurgenson, S., Soucy, E. R., Kim, H. S., & Reid, R. C. (2011). Network anatomy and in vivo physiology of visual cortical neurons. Nature, 471(7337), 177–182. 10.1038/nature09802

Bosch, M., & Hayashi, Y. (2012). Structural plasticity of dendritic spines. Current opinion in neurobiology, 22(3), 383–388. 10.1016/j.conb.2011.09.002

Bourne, J. N., & Harris, K. M. (2008). Balancing structure and function at hippocampal dendritic spines. Annual review of neuroscience, 31, 47–67. 10.1146/annurev.neuro.31.060407.125646

Campello, R.J.G.B., Moulavi, D., Sander, J. (2013). Density-Based Clustering Based on Hierarchical Density Estimates. In: Pei, J., Tseng, V.S., Cao, L., Motoda, H., Xu, G. (eds) Advances in Knowledge Discovery and Data Mining. PAKDD 2013. Lecture Notes in Computer Science(), vol 7819. Springer, Berlin, Heidelberg. 10.1007/978-3-642-37456-2_14

Cano-Astorga, N., DeFelipe, J., & Alonso-Nanclares, L. (2021). Three-Dimensional Synaptic Organization of Layer III of the Human Temporal Neocortex. Cerebral cortex (New York, N.Y. : 1991), 31(10), 4742–4764. 10.1093/cercor/bhab120

Cano-Astorga, N., Plaza-Alonso, S., DeFelipe, J., & Alonso-Nanclares, L. (2024). Volume electron microscopy analysis of synapses in primary regions of the human cerebral cortex. Cerebral cortex (New York, N.Y. : 1991), 34(8), bhae312. 10.1093/cercor/bhae312

Colgan, L. A., Parra-Bueno, P., Holman, H. L., Tu, X., Jain, A., Calubag, M. F., Misler, J. A., Gary, C., Oz, G., Suponitsky-Kroyter, I., Okaz, E., & Yasuda, R. (2023). Dual Regulation of Spine-Specific and Synapse-to-Nucleus Signaling by PKCδ during Plasticity. The Journal of neuroscience : the official journal of the Society for Neuroscience, 43(30), 5432–5447. 10.1523/JNEUROSCI.0208-22.2023

Cornejo, V. H., Ofer, N., & Yuste, R. (2022). Voltage compartmentalization in dendritic spines in vivo. Science (New York, N.Y.), 375(6576), 82–86. 10.1126/science.abg0501

Del Giudice M. (2021). Effective Dimensionality: A Tutorial. Multivariate behavioral research, 56(3), 527–542. 10.1080/00273171.2020.1743631

Domínguez-Álvaro, M., Montero-Crespo, M., Blazquez-Llorca, L., Plaza-Alonso, S., Cano-Astorga, N., DeFelipe, J., & Alonso-Nanclares, L. (2021). 3D Analysis of the Synaptic Organization in the Entorhinal Cortex in Alzheimer’s Disease. eNeuro, 8(3), ENEURO.0504-20.2021. 10.1523/ENEURO.0504-20.2021

Fernholz, M. H. P., Guggiana Nilo, D. A., Bonhoeffer, T., & Kist, A. M. (2024). DeepD3, an open framework for automated quantification of dendritic spines. PLoS computational biology, 20(2), e1011774. 10.1371/journal.pcbi.1011774

Fish, K. N., Sweet, R. A., MacDonald, M. L., & Lewis, D. A. (2025). Regional Specificity of Cortical Layer 3 Dendritic Spine Deficits in Schizophrenia. JAMA psychiatry, 82(11), 1123–1132. 10.1001/jamapsychiatry.2025.2221

Glantz, L. A., & Lewis, D. A. (2000). Decreased dendritic spine density on prefrontal cortical pyramidal neurons in schizophrenia. Archives of general psychiatry, 57(1), 65–73. 10.1001/archpsyc.57.1.65

Glausier, J. R., & Lewis, D. A. (2013). Dendritic spine pathology in schizophrenia. Neuroscience, 251, 90–107. 10.1016/j.neuroscience.2012.04.044

Grutzendler, J., Kasthuri, N., & Gan, W. B. (2002). Long-term dendritic spine stability in the adult cortex. Nature, 420(6917), 812–816. 10.1038/nature01276

Harris, C. R., Millman, K. J., van der Walt, S. J., Gommers, R., Virtanen, P., Cournapeau, D., Wieser, E., Taylor, J., Berg, S., Smith, N. J., Kern, R., Picus, M., Hoyer, S., van Kerkwijk, M. H., Brett, M., Haldane, A., Del Río, J. F., Wiebe, M., Peterson, P., Gérard-Marchant, P., … Oliphant, T. E. (2020). Array programming with NumPy. Nature, 585(7825), 357–362. 10.1038/s41586-020-2649-2

Harris, K. M., Jensen, F. E., & Tsao, B. (1992). Three-dimensional structure of dendritic spines and synapses in rat hippocampus (CA1) at postnatal day 15 and adult ages: implications for the maturation of synaptic physiology and long-term potentiation. The Journal of neuroscience : the official journal of the Society for Neuroscience, 12(7), 2685–2705. 10.1523/JNEUROSCI.12-07-02685.1992

Harris, K. M., Kuwajima, M., Flores, J. C., & Zito, K. (2024). Synapse-specific structural plasticity that protects and refines local circuits during LTP and LTD. Philosophical transactions of the Royal Society of London. Series B, Biological sciences, 379(1906), 20230224. 10.1098/rstb.2023.0224

Harris, K. M., & Kater, S. B. (1994). Dendritic spines: cellular specializations imparting both stability and flexibility to synaptic function. Annual review of neuroscience, 17, 341–371. 10.1146/annurev.ne.17.030194.002013

Harris, K. M., & Stevens, J. K. (1989). Dendritic spines of CA 1 pyramidal cells in the rat hippocampus: serial electron microscopy with reference to their biophysical characteristics. The Journal of neuroscience : the official journal of the Society for Neuroscience, 9(8), 2982–2997. 10.1523/JNEUROSCI.09-08-02982.1989

Harvey, C. D., & Svoboda, K. (2007). Locally dynamic synaptic learning rules in pyramidal neuron dendrites. Nature, 450(7173), 1195–1200. 10.1038/nature06416

Holtmaat, A., & Svoboda, K. (2009). Experience-dependent structural synaptic plasticity in the mammalian brain. Nature reviews. Neuroscience, 10(9), 647–658. 10.1038/nrn2699

Hotulainen, P., & Hoogenraad, C. C. (2010). Actin in dendritic spines: connecting dynamics to function. The Journal of cell biology, 189(4), 619–629. 10.1083/jcb.201003008

Jacot-Descombes, S., Keshav, N. U., Dickstein, D. L., Wicinski, B., Janssen, W. G. M., Hiester, L. L., Sarfo, E. K., Warda, T., Fam, M. M., Harony-Nicolas, H., Buxbaum, J. D., Hof, P. R., & Varghese, M. (2020). Altered synaptic ultrastructure in the prefrontal cortex of Shank3-deficient rats. Molecular autism, 11(1), 89. 10.1186/s13229-020-00393-8

Jain, A., Nakahata, Y., Pancani, T., Watabe, T., Rusina, P., South, K., Adachi, K., Yan, L., Simorowski, N., Furukawa, H., & Yasuda, R. (2024). Dendritic, delayed, stochastic CaMKII activation in behavioural time scale plasticity. Nature, 635(8037), 151–159. 10.1038/s41586-024-08021-8

Januszewski, M., Kornfeld, J., Li, P. H., Pope, A., Blakely, T., Lindsey, L., Maitin-Shepard, J., Tyka, M., Denk, W., & Jain, V. (2018). High-precision automated reconstruction of neurons with flood-filling networks. Nature methods, 15(8), 605–610. 10.1038/s41592-018-0049-4

Kasai, H., Fukuda, M., Watanabe, S., Hayashi-Takagi, A., & Noguchi, J. (2010). Structural dynamics of dendritic spines in memory and cognition. Trends in neurosciences, 33(3), 121–129. 10.1016/j.tins.2010.01.001

Kasai, H., Ucar, H., Morimoto, Y., Eto, F., & Okazaki, H. (2023). Mechanical transmission at spine synapses: Short-term potentiation and working memory. Current opinion in neurobiology, 80, 102706. 10.1016/j.conb.2023.102706

Kashiwagi, Y., Higashi, T., Obashi, K., Sato, Y., Komiyama, N. H., Grant, S. G. N., & Okabe, S. (2019). Computational geometry analysis of dendritic spines by structured illumination microscopy. Nature communications, 10(1), 1285. 10.1038/s41467-019-09337-0

Kasthuri, N., Hayworth, K. J., Berger, D. R., Schalek, R. L., Conchello, J. A., Knowles-Barley, S., Lee, D., Vázquez-Reina, A., Kaynig, V., Jones, T. R., Roberts, M., Morgan, J. L., Tapia, J. C., Seung, H. S., Roncal, W. G., Vogelstein, J. T., Burns, R., Sussman, D. L., Priebe, C. E., Pfister, H., … Lichtman, J. W. (2015). Saturated Reconstruction of a Volume of Neocortex. Cell, 162(3), 648–661. 10.1016/j.cell.2015.06.054

Koch, C., & Zador, A. (1993). The function of dendritic spines: devices subserving biochemical rather than electrical compartmentalization. The Journal of neuroscience : the official journal of the Society for Neuroscience, 13(2), 413–422. 10.1523/JNEUROSCI.13-02-00413.1993

Larkum, M. E., Zhu, J. J., & Sakmann, B. (1999). A new cellular mechanism for coupling inputs arriving at different cortical layers. Nature, 398(6725), 338–341. 10.1038/18686

Levet, F., Tønnesen, J., Nägerl, U. V., & Sibarita, J. B. (2020). SpineJ: A software tool for quantitative analysis of nanoscale spine morphology. Methods (San Diego, Calif.), 174, 49–55. 10.1016/j.ymeth.2020.01.020

Lichtman, J. W., & Denk, W. (2011). The big and the small: challenges of imaging the brain’s circuits. Science (New York, N.Y.), 334(6056), 618–623. 10.1126/science.1209168

Lisman, J. E., & Harris, K. M. (1993). Quantal analysis and synaptic anatomy--integrating two views of hippocampal plasticity. Trends in neurosciences, 16(4), 141–147. 10.1016/0166-2236(93)90122-3

Loomba, S., Straehle, J., Gangadharan, V., Heike, N., Khalifa, A., Motta, A., Ju, N., Sievers, M., Gempt, J., Meyer, H. S., & Helmstaedter, M. (2022). Connectomic comparison of mouse and human cortex. Science (New York, N.Y.), 377(6602), eabo0924. 10.1126/science.abo0924

Luengo-Sanchez, S., Fernaud-Espinosa, I., Bielza, C., Benavides-Piccione, R., Larrañaga, P., & DeFelipe, J. (2018). 3D morphology-based clustering and simulation of human pyramidal cell dendritic spines. PLoS computational biology, 14(6), e1006221. 10.1371/journal.pcbi.1006221

Manjarrez, E., Hernández, S.T., Zamora-Ursulo, M.A., & Flores, A. (2026). Rotating a petavoxel reconstruction exposes the viewing-angle bias inherent to Golgi-Cox and confocal dendritic-spine classification. BioRxiv. 10.64898/2026.06.15.732500

Matsuzaki, M., Ellis-Davies, G. C., Nemoto, T., Miyashita, Y., Iino, M., & Kasai, H. (2001). Dendritic spine geometry is critical for AMPA receptor expression in hippocampal CA1 pyramidal neurons. Nature neuroscience, 4(11), 1086–1092. 10.1038/nn736

Matsuzaki, M., Honkura, N., Ellis-Davies, G. C., & Kasai, H. (2004). Structural basis of long-term potentiation in single dendritic spines. Nature, 429(6993), 761–766. 10.1038/nature02617

McInnes, L., Healy, J., & Astels, S. (2017). HDBSCAN: Hierarchical density based clustering. Journal of Open Source Software, 2, 205. 10.21105/joss.00205

McKinney, W. (2010). Data structures for statistical computing in Python. Proceedings of the 9th Python in Science Conference, 51–56. 10.25080/Majora-92bf1922-00a

Montero-Crespo, M., Domínguez-Álvaro, M., Alonso-Nanclares, L., DeFelipe, J., & Blazquez-Llorca, L. (2021). Three-dimensional analysis of synaptic organization in the hippocampal CA1 field in Alzheimer’s disease. Brain : a journal of neurology, 144(2), 553–573. 10.1093/brain/awaa406

Montero-Crespo, M., Dominguez-Alvaro, M., Rondon-Carrillo, P., Alonso-Nanclares, L., DeFelipe, J., & Blazquez-Llorca, L. (2020). Three-dimensional synaptic organization of the human hippocampal CA1 field. eLife, 9, e57013. 10.7554/eLife.57013

Motta, A., Berning, M., Boergens, K. M., Staffler, B., Beining, M., Loomba, S., Hennig, P., Wissler, H., & Helmstaedter, M. (2019). Dense connectomic reconstruction in layer 4 of the somatosensory cortex. Science (New York, N.Y.), 366(6469), eaay3134. 10.1126/science.aay3134

Nicoll, R. A., & Schulman, H. (2023). Synaptic memory and CaMKII. Physiological reviews, 103(4), 2877–2925. 10.1152/physrev.00034.2022

Nimchinsky, E. A., Sabatini, B. L., & Svoboda, K. (2002). Structure and function of dendritic spines. Annual review of physiology, 64, 313–353. 10.1146/annurev.physiol.64.081501.160008

Ofer, N., Benavides-Piccione, R., DeFelipe, J., & Yuste, R. (2022). Structural Analysis of Human and Mouse Dendritic Spines Reveals a Morphological Continuum and Differences across Ages and Species. eNeuro, 9(3), ENEURO.0039-22.2022. 10.1523/ENEURO.0039-22.2022

Ofer, N., Berger, D. R., Kasthuri, N., Lichtman, J. W., & Yuste, R. (2021). Ultrastructural analysis of dendritic spine necks reveals a continuum of spine morphologies. Developmental neurobiology, 81(5), 746–757. 10.1002/dneu.22829

Pchitskaya, E., & Bezprozvanny, I. (2020). Dendritic Spines Shape Analysis-Classification or Clusterization? Perspective. Frontiers in synaptic neuroscience, 12, 31. 10.3389/fnsyn.2020.00031

Pedregosa, F., Varoquaux, G., Gramfort, A., Michel, V., Thirion, B., Grisel, O., Blondel, M., Prettenhofer, P., Weiss, R., Dubourg, V., Vanderplas, J. & others. (2011). Scikit-learn: Machine learning in Python. Journal of Machine Learning Research, 12, 2825–2830. https://www.jmlr.org/papers/volume12/pedregosa11a/pedregosa11a.pdf

Penzes, P., Cahill, M. E., Jones, K. A., VanLeeuwen, J. E., & Woolfrey, K. M. (2011). Dendritic spine pathology in neuropsychiatric disorders. Nature neuroscience, 14(3), 285–293. 10.1038/nn.2741

Peters, A., & Kaiserman-Abramof, I. R. (1970). The small pyramidal neuron of the rat cerebral cortex. The perikaryon, dendrites and spines. The American journal of anatomy, 127(4), 321–355. 10.1002/aja.1001270402

Shapson-Coe, A., Januszewski, M., Berger, D. R., Pope, A., Wu, Y., Blakely, T., Schalek, R. L., Li, P. H., Wang, S., Maitin-Shepard, J., Karlupia, N., Dorkenwald, S., Sjostedt, E., Leavitt, L., Lee, D., Troidl, J., Collman, F., Bailey, L., Fitzmaurice, A., Kar, R., … Lichtman, J. W. (2024). A petavoxel fragment of human cerebral cortex reconstructed at nanoscale resolution. Science (New York, N.Y.), 384(6696), eadk4858. 10.1126/science.adk4858

Spruston N. (2008). Pyramidal neurons: dendritic structure and synaptic integration. Nature reviews. Neuroscience, 9(3), 206–221. 10.1038/nrn2286

Steffens, H., Mott, A. C., Li, S., Wegner, W., Švehla, P., Kan, V. W. Y., Wolf, F., Liebscher, S., & Willig, K. I. (2021). Stable but not rigid: Chronic in vivo STED nanoscopy reveals extensive remodeling of spines, indicating multiple drivers of plasticity. Science advances, 7(24), eabf2806. 10.1126/sciadv.abf2806

Südhof T. C. (2008). Neuroligins and neurexins link synaptic function to cognitive disease. Nature, 455(7215), 903–911. 10.1038/nature07456

Sullivan, C. B., & Kaszynski, A. (2019). PyVista: 3D plotting and mesh analysis through a streamlined interface for the Visualization Toolkit (VTK). Journal of Open Source Software, 4, 1450. 10.21105/joss.01450

Taubin, G. (1995). A signal processing approach to fair surface design. Proceedings of the 22nd Annual Conference on Computer Graphics and Interactive Techniques (SIGGRAPH ’95), 351–358. 10.1145/218380.218473

Terry, R. D., Masliah, E., Salmon, D. P., Butters, N., DeTeresa, R., Hill, R., Hansen, L. A., & Katzman, R. (1991). Physical basis of cognitive alterations in Alzheimer’s disease: synapse loss is the major correlate of cognitive impairment. Annals of neurology, 30(4), 572–580. 10.1002/ana.410300410

Tønnesen, J., Katona, G., Rózsa, B., & Nägerl, U. V. (2014). Spine neck plasticity regulates compartmentalization of synapses. Nature neuroscience, 17(5), 678–685. 10.1038/nn.3682

Trachtenberg, J. T., Chen, B. E., Knott, G. W., Feng, G., Sanes, J. R., Welker, E., & Svoboda, K. (2002). Long-term in vivo imaging of experience-dependent synaptic plasticity in adult cortex. Nature, 420(6917), 788–794. 10.1038/nature01273

Turner, N. L., Macrina, T., Bae, J. A., Yang, R., Wilson, A. M., Schneider-Mizell, C., Lee, K., Lu, R., Wu, J., Bodor, A. L., Bleckert, A. A., Brittain, D., Froudarakis, E., Dorkenwald, S., Collman, F., Kemnitz, N., Ih, D., Silversmith, W. M., Zung, J., Zlateski, A., … Seung, H. S. (2022). Reconstruction of neocortex: Organelles, compartments, cells, circuits, and activity. Cell, 185(6), 1082–1100.e24. 10.1016/j.cell.2022.01.023

Van Rossum, G., & Drake, F. L. (2009). Python 3 Reference Manual. CreateSpace.100 Enterprise Way, Suite A200Scotts ValleyCA

Virtanen, P., Gommers, R., Oliphant, T. E., Haberland, M., Reddy, T., Cournapeau, D., Burovski, E., Peterson, P., Weckesser, W., Bright, J., van der Walt, S. J., Brett, M., Wilson, J., Millman, K. J., Mayorov, N., Nelson, A. R. J., Jones, E., Kern, R., Larson, E., Carey, C. J., … SciPy 1.0 Contributors (2020). SciPy 1.0: fundamental algorithms for scientific computing in Python. Nature methods, 17(3), 261–272. 10.1038/s41592-019-0686-2

Walker, C. K., Liu, E., Greathouse, K. M., Adamson, A. B., Wilson, J. P., Poovey, E. H., Curtis, K. A., Muhammad, H. M., Weber, A. J., Bennett, D. A., Seyfried, N. T., Gaiteri, C., & Herskowitz, J. H. (2024). Dendritic spine head diameter predicts episodic memory performance in older adults. Science advances, 10(32), eadn5181. 10.1126/sciadv.adn5181

Yang, Y., & Liu, J. J. (2022). Structural LTP: Signal transduction, actin cytoskeleton reorganization, and membrane remodeling of dendritic spines. Current opinion in neurobiology, 74, 102534. 10.1016/j.conb.2022.102534

Yasuda, R., Hayashi, Y., & Hell, J. W. (2022). CaMKII: a central molecular organizer of synaptic plasticity, learning and memory. Nature reviews. Neuroscience, 23(11), 666–682. 10.1038/s41583-022-00624-2

Yuste R. (2011). Dendritic spines and distributed circuits. Neuron, 71(5), 772–781. 10.1016/j.neuron.2011.07.024

Yuste R. (2013). Electrical compartmentalization in dendritic spines. Annual review of neuroscience, 36, 429–449. 10.1146/annurev-neuro-062111-150455

Yuste, R., & Bonhoeffer, T. (2001). Morphological changes in dendritic spines associated with long-term synaptic plasticity. Annual review of neuroscience, 24, 1071–1089. 10.1146/annurev.neuro.24.1.1071

Yuste, R., & Bonhoeffer, T. (2004). Genesis of dendritic spines: insights from ultrastructural and imaging studies. Nature reviews. Neuroscience, 5(1), 24–34. 10.1038/nrn1300

